# Porous microneedle patch with sustained exosome delivery repairs severe spinal cord injury

**DOI:** 10.1101/2022.07.18.500400

**Authors:** Ao Fang, Yifan Wang, Naiyu Guan, Lingmin Lin, Binjie Guo, Wanxiong Cai, Xiangfeng Chen, Jingjia Ye, Zeinab Abdelrahman, Xiaodan Li, Yanming Zuo, Hanyu Zheng, Zhonghan Wu, Shuang Jin, Kan Xu, Xiaosong Gu, Bin Yu, Xuhua Wang

## Abstract

Mesenchymal stem cell-derived exosome (MSC-EXO) transplantation has been suggested as an efficacious treatment to suppress spinal cord injury (SCI)-triggered neuroinflammation. However, an ethically acceptable method to continuously deliver MSC-EXOs to acute spinal lesions, without damaging nearby tissues/axons, has never been achieved. In this study, we fabricated a device comprising a patch containing MSCs and a microneedle array (MN-MSC patch) to treat severe SCI. When topically applied to an acute spinal lesion beneath the spinal dura, the soft microneedle (MN) array with reasonable mechanical strength avoided damaging the nearby spinal tissues, and the porous microstructure of MNs facilitated highly efficient MSC-EXO delivery. With the capacity for sustained delivery of MSC-EXOs, the MN-MSC patch was evaluated in a contusive rat SCI model. The MSCs encapsulated in the patch could survive for at least 7 days, encompassing the optimal time window for downregulating SCI-triggered neuroinflammation. As a result, MN-MSC patch treatment led to reduced cavity and scar tissue formation, greater angiogenesis, and improved survival of nearby tissues/axons. Remarkably, rats treated by this method achieved superior muscle control and exhibited robust hindlimb locomotion functional recovery. Conclusively, the MN-MSC patch device proposed here overcomes the current dilemma between treatment efficacy and ethical issues in treating acute SCI.

## 2. Introduction

There are approximately 930,000 new spinal cord injury (SCI) cases caused by accidents each year^1^. Even though the systemic administration of methylprednisolone sodium succinate (MPSS) has been approved as the only drug for treating acute SCI, this drug has limited therapeutic efficacy and severe side effects, which severely limits its application in clinical practice^2–4^. SCI is initially caused by mechanical trauma and diverse mechanisms of secondary damage following the inflammatory response^5^. Secondary damage can cause hemorrhage, edema, decreased blood flow and inflammation, thus affecting adjacent spinal cord segments^6^. The inflammatory environment triggered by the activation of microglia and the release of proinflammatory cytokines leads to inevitable neuronal damage and, consequently, scar and cystic cavity formation in spinal lesion sites^7,8^. Recently, the transplantation of mesenchymal stem cells (MSCs) into lesion sites has been suggested as an effective treatment to alleviate the neuroinflammation triggered by SCI^9^ and promote healing after injury^10–14^. However, the neuroinflammatory microenvironment of SCI lesions is harmful to MSC survival, significantly impeding the clinical application of these cells^15^.

Fortunately, MSC-derived exosomes (MSC-EXOs) secreted by MSCs have similar therapeutic benefits for immune modulation when injected into acute SCI lesions^16–20^. Previous studies have shown that MSC-EXOs immobilized in a peptide-modified adhesive hydrogel transplanted into an SCI lesion during the acute phase could mitigate the SCI microenvironment and promote functional recovery^21,22^. However, the retention time of MSC-EXOs in spinal tissue might not be long enough for them to exhibit their optimal therapeutic effects. The neuroinflammation and neuronal apoptosis caused by SCI could continue for a couple of weeks or even longer^23,24^, but MSC-EXOs could not be continuously delivered to SCI lesions by these methods. Moreover, direct transplantation of MSC-EXOs into the spinal injury site might damage the nearby healthy tissue adjacent to the lesion, cause unpredictable side effects and increase medical accidents^25–31^. Therefore, in clinical practice, the direct transplantation of MSC-EXOs or materials containing MSC-EXOs into the spinal lesion of an acute SCI patient is not suited for ethical approval^32^.

To overcome these challenges, we took advantage of a microneedle (MN) array, which has facilitated painless localized delivery of drugs or therapeutic biomolecules with good tolerability in clinical trials^33–36^. We then fabricated a patch with an MN array with good mechanical strength matching soft spinal tissues and with a suitable pore size for MSC-EXO delivery when it was mounted onto a spinal cord lesion beneath the dura. To sustainably deliver MSC-EXOs, the MN arrays were additionally mounted with a gelatin methacryloyl (GelMA) hydrogel block embedded with MSCs during surgery (Scheme 1). We envisioned that the GelMA hydrogel block could provide a biocompatible microenvironment for the long-term survival of MSCs that can secrete and achieve sustained delivery of MSC-EXOs to the spinal lesion for a long time. In this way, the MSC-EXOs can be efficiently and precisely delivered to SCI lesions through the porous MNs for optimal therapeutic efficacy without additional damage (Scheme 1). We showed that the MN-MSC patch could remain on the spinal injury sites and sustainably release MSC-EXOs for at least one week, which resulted in remarkable tissue repair, superior muscle control and robust functional recovery after severe SCI (Scheme 1).

## 3. Results

### 3.1 Fabrication and Characterization of the Patch with an MN array *In Vitro*

To avoid injecting MSCs into the spinal cord and damaging the healthy spinal tissue adjacent to the spinal cord lesions, we designed a patch with an MN array (Fig. 1a). The MN patch was made from chemically cross-linked methacryloyl (MW. 100-250 kDa) with a micromolding method to form GelMA (Fig. 1a, a1, a2)^35^. The scale of the fabricated MN patch was approximately 4 mm × 4 mm with 45 needles, which is large enough to cover a typical rat contusive spinal cord lesion. The needle of the MN arrays was conical, with a base diameter of 250 μm, a height of 600 μm and a tip diameter of 10 μm (Fig. 1e-g). To sustainably release exosomes, the patch was additionally covered with GelMA hydrogel embedded with MSCs during surgery (Supplementary Fig. S2d-g). The GelMA hydrogel with porous structures was designed to facilitate the release of MSC-EXOs through the polymeric needles. As shown by the scanning electron microscopy (SEM) images, the pore size of the gel reached up to approximately 100 μm (Fig. 1b, b2, b4), compared with the relatively small pore size of 10 μm in the traditional GelMA hydrogel (Fig. 1b, b1, b3).

**Figure 1.**
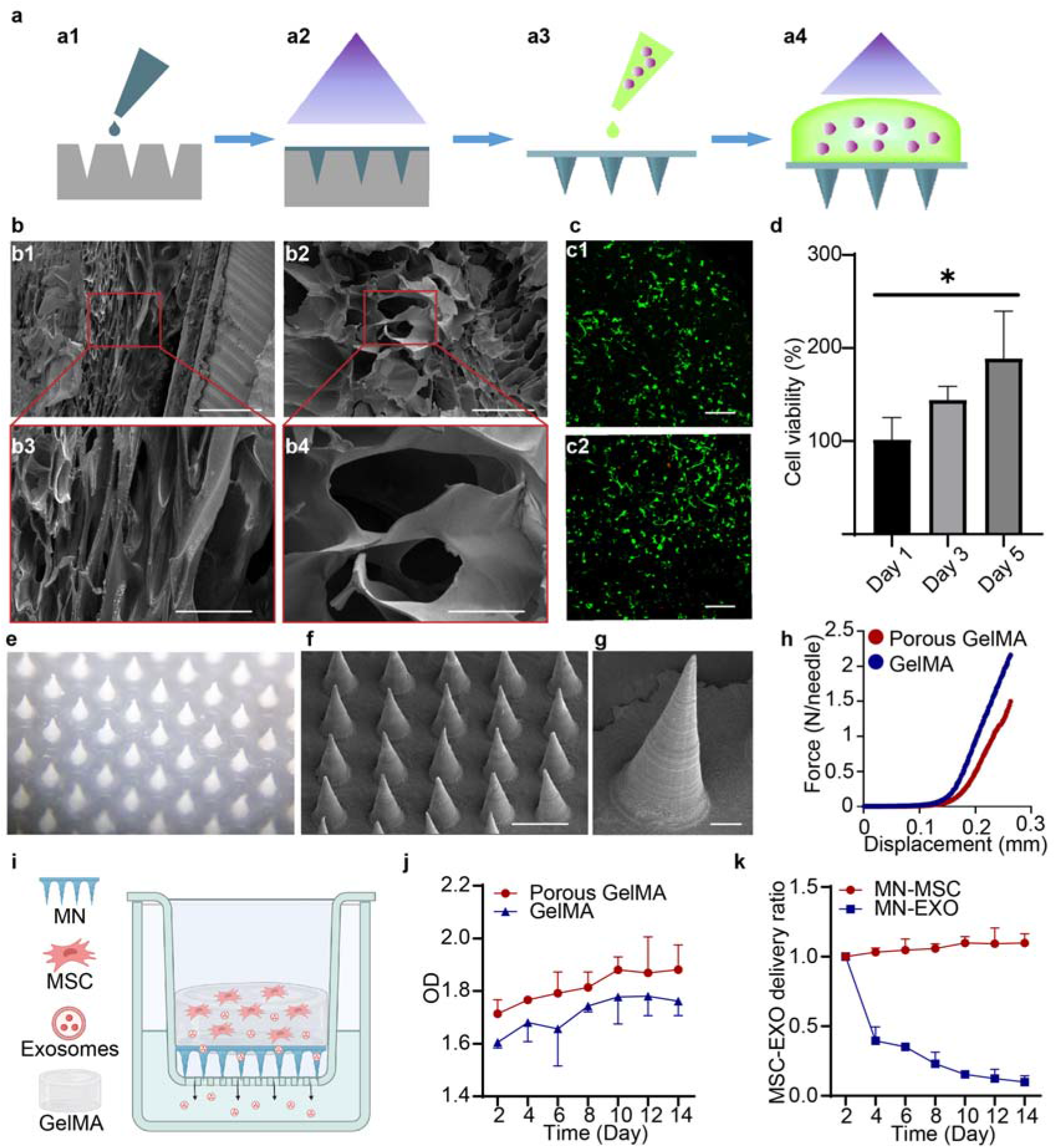
Fabrication and characterization of the MN-MSC patch. a) Schematic of the MN-MSC patch fabrication process by a1) casting, a2) blue light crosslinking, a3) peeling from the PDMS mold and adding GelMA solution with MSCs, and a4) blue light crosslinking. b) SEM image of the b1) interstructure of normal and b2) porous GelMA hydrogels. Scale bars of images (b1, b2) indicate 300 μm, and scale bars of images (b3, b4) indicate 100 μm. c) Calcein (live)/EthD (dead) staining revealed the morphology and viability of MSCs encapsulated in GelMA hydrogel on c1) day 3 and c2) day 5. Scale bars: 200 μm. d) Quantitative viability analysis of MSCs encapsulated in GelMA hydrogel on days 1, 3 and 5. Data were analyzed with one-way ANOVA followed by Tukey’s post-hoc test. n=4. *p<0.05. e) Representative optical microscopy images and f-g) SEM images of MNs. Scale bars for images f) indicate 500 μm, and scale bars for images g) indicate 50 μm. h) The mechanical strength of MNs. i) Schematic showing the study design used to test the release profile of exosomes from the MN-MSC patch. j) The daily exosome release curves of MN-MSC patches fabricated with porous and normal GelMA hydrogels. Data are presented as the means ± SDs (n = 3 for each group). k) The daily exosome release curves of MN-MSC and MN-EXO patches. Data are presented as the means ± SDs (n = 3 for each group).

To examine the biocompatibility of the MN patch, we first dropped the methacryloyl solution with MSCs (approximately 2.5×10^7^ cells/mL) onto the surface of the patch and cross-linked methacryloyl to form GelMA hydrogel with blue light irradiation (Fig. 1a, a3, a4). The live/dead staining of the GelMA hydrogel revealed excellent viability of embedded MSCs after culturing for 3 and 5 days (Fig. 1c). Quantitative analysis with the CCK-8 test indicated that MSCs in the GelMA hydrogel of the MN-MSC patch survived and proliferated well (Fig. 1d). The mechanical strength of each needle constructed by the porous structure matched that of the soft spinal cord tissue^37^. The needle with the porous structure had a relatively low mechanical strength compared to that of the normal one, which is propitious for preventing spinal cord damage (Fig. 1h). We next assessed the MSC-EXO release capacity of the MN patch with this porous structure. To do so, the patch made from porous GelMA^38^ or regular GelMA was seeded with MSCs and placed in the top well of a transwell (Fig. 1i). The transwells were then immersed in culture media and incubated at 37 °C. At regular intervals, aliquots of samples were collected from the bottom wells, and the concentrations of MSC-EXOs in the samples were measured by a Micro BCA Protein Assay kit^39^. Plotting the optical density (OD) of the samples revealed that the MN patch made from porous GelMA released MSC-EXOs more efficiently (Fig. 1j), possibly because the MN patch with a larger pore size was more permissive for MSC-EXOs. We therefore chose the MN patch made from porous GelMA for the following studies.

Previous studies have shown that most of the therapeutic benefits of MSCs in SCI treatment are mediated by MSC-EXO release^40^, highlighting the importance of sustained MSC-EXO delivery in improving therapeutic efficacy. We therefore sought to evaluate the sustained MSC-EXO delivery capacity of MN patches embedded with MSCs (MN-MSCs), and MN patches loaded with MSC-EXOs (MN-EXOs) were used as controls. The MSC-EXOs were isolated from MSC culture supernatant via ultracentrifugation and showed a typical cup-shaped morphology (Supplementary Fig. S1a). Through nanoparticle tracking analysis (NTA), we found that the MSC-EXOs had an average size of 110 nm, which was consistent with the results of previous studies (Supplementary Fig. S1c)^41^. The expression of tetraspanins CD9 and TSG101 (mesenchymal markers) on the collected samples (Supplementary Fig. S1d) indicated successful isolation and purification of MSC-EXOs. We then compared the persistent MSC-EXO release capacity of MSC-seeded and MSC-EXO-loaded MN patches. Plotting the relative concentration of released MSC-EXOs in the collected samples from the bottom wells of the transwells at different time points revealed that the release of MSC-EXOs from MSC-EXO-loaded MN patches quickly decreased in the first 4 days, whereas the delivery of MSC-EXOs from MSC-seeded MN patches was stable but with a slight increase at 2 weeks (Fig. 1k), indicating that the MN patches had good biocompatibility for MSC survival. Thus, we fabricated a device to persistently deliver MSC-EXOs with high efficiency.

### 3.2 Distribution of Exosomes in the Injured Spinal Cord after MN Patch Implantation

After verifying that the fabricated MN-MSC patch could sustainably deliver exosomes *in vitro*, we sought to assess whether the MSC-EXOs of the patch could be delivered through the MNs into injured spinal cord tissues in a contusive rat SCI model. We used a severe T10 contusive SCI model, which was constructed by an infinite vertical impactor with the parameters of a 2.5 m/s rate, 2 mm depth, 5 s duration time and 3 mm diameter cylinder^42^. Because MSC-EXOs in live MSCs are difficult to label, MSC-EXOs stained with CM-DiI were encapsulated into GelMA hydrogels in this experiment. The solid red fluorescence of MSC-EXOs encapsulated in the GelMA hydrogel indicated their successful labeling with CM-DiI (Supplementary Fig. S1b). After contusive SCI, the dura of the spinal cord was removed carefully by a pair of surgical scissors. Then, the MN patch was placed on top of the spinal lesion, and a solution of GelMA mixed with MSC-EXOs was mounted on the top of the MN patch, followed by a blue light photocuring process (Fig. 2a). Seven days after surgery, we found that most of the released MSC-EXOs were localized and aggregated in the spinal injury site (Fig. 2b and c, b1, b2, c1, c2), indicating the localized delivery capacity of the MN patch. Conclusively, the MN patch has excellent localized delivery capacity of MSC-EXOs *in vivo*, which facilitates sustained MSC-EXO delivery for SCI treatment.

**Figure 2.**
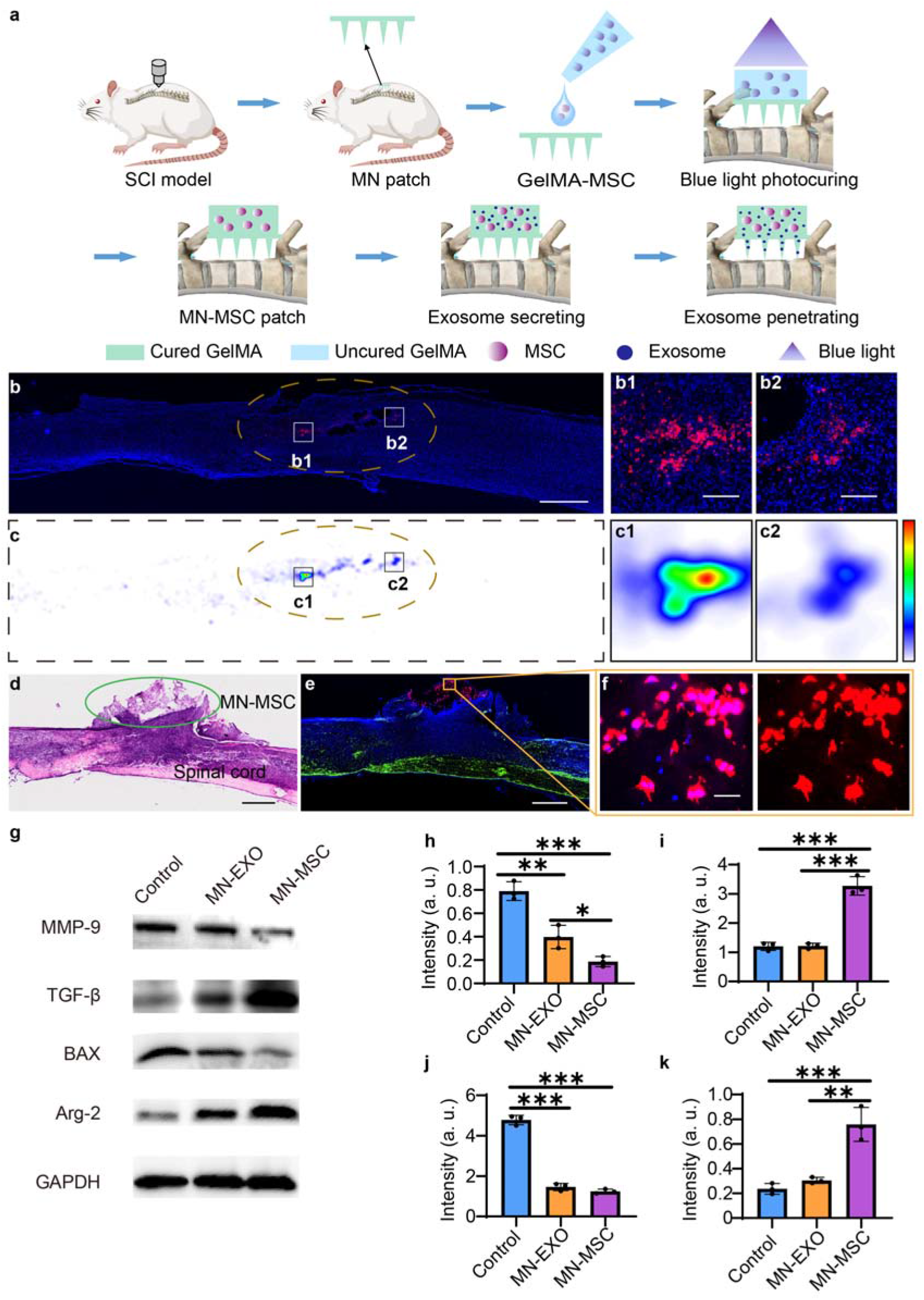
Sustained MSC-EXO delivery by MN-MSC patch treatment alleviates neuroinflammation triggered by SCI. a) Schematic illustration of MN-MSC patch implantation on the injury site of the spinal cord. b) *In vivo* distribution of exosomes (DiI labeled, red) in the injured spinal cord tissues after MN-EXO patch implantation, blue (DAPI). Scale bar: 1 mm. The injury region of the spinal cord is marked with yellow dotted lines. c) Heatmaps of exosome distribution in the injured spinal cord tissues after MN-EXO patch implantation; red: the highest numbers of exosomes, blue: the lowest, and white: background. The injury region of the spinal cord is marked with yellow dotted lines. b1, b2, c1, c2) Enlarged images to show the details. Scale bar: 100 μm. d) HE staining of the injured spinal cord with MN-MSC patch implantation; scale bar: 100 μm. e) Representative images of immunohistochemical staining for MN-MSC patches on the injured spinal cord on day 7 after SCI. Green (GFAP), blue (DAPI), red (GAPDH), scale bar: 1 mm. f) High-magnification images of the boxed area, scale bar: 50 μm. g-k) WB results of quantitative analyses of h) MMP-9, i) TGF-β, j) BAX and k) Arg-2 expression in the first week at the injury site of rats in the control, MN-EXO, and MN-MSC groups. Statistical analysis was performed using one-way ANOVA followed by Tukey’s multiple comparisons test. n=3. **p<0.01, ***p<0.001.

Next, we attempted to assess whether the cells in the MN-MSC patch could survive long enough to ameliorate the SCI-induced neuroinflammatory environment and protect the spared tissues/axons from secondary injuries. We implanted the MN patch into the spinal lesion of the same SCI model, and the solution of MSCs mixed with GelMA was mounted on the top of the MN patch before the blue light photocuring process was performed (Fig. 2a, S2a-h). As we know, the optimal time for suppressing acute SCI-triggered neuroinflammation and protecting the spared tissue/axons is within 7 days after SCI^42^, so we dissected the spinal cord on the 7^th^ day after SCI to assess whether MSCs could survive in the MN-MSC patch until this time point. As indicated by the images of spinal sections stained with hematoxylin and eosin (H&E) (Fig. 2d), the MN patch still persisted on the spinal cord surface at this time. More importantly, staining of the human MSC marker GAPDH in the spinal tissues indicated that the MSCs had survived well in the MN-MSC patch on the 7^th^ day after SCI (Fig. 2e, f, red). These results confirmed that the MSCs in the MN-MSC patch can survive at the injury site long enough to cover the optimal therapeutic time window of SCI. Moreover, immunofluorescent staining of a serial spinal section from the spinal border to the center showed that MSCs from the MN-MSC patch did not migrate from the GelMA hydrogel block into the spinal cord (Fig. S3). These results indicated that the fabricated device achieved our aim to sustainably deliver MSC-EXOs without MSC invasion into the spinal cord.

### 3.3 The MN-MSC Patch Alleviates the Inflammatory Response in Spinal Lesions after SCI

To assess whether the MN-MSC patch could the SCI-induced neuroinflammatory environment, we constructed animal models without any treatment (control group), with MN patches embedded with MSC-EXO implantation (MN-EXO group) or with MN patches seeded with MSC implantation (MN-MSC group) for comparison. On day 7, we performed western blotting analysis to detect the protein expression levels of the spinal tissues with spinal lesions in rats that underwent different treatments. We found that the expression of the proinflammatory MMP-9 marker was significantly reduced and the expression of the anti-inflammatory TGF-β marker was significantly increased in the spinal tissues of rats treated with the MN-MSC patch (Fig. 2g, h, i). In addition, the expression of the proapoptotic marker BAX was significantly reduced (Fig. 2g, j), and the angiogenic Arg-2 marker was significantly increased in the spinal tissues of rats that had the MN-MSC patch treatments (Fig. 2g, k). In contrast, we found nearly no significant differences in the expression of these genes between the rats in the MN-EXO and control groups (Fig. 2g-k), possibly because the limited number of exosomes delivered by MN-EXOs was not enough to significantly suppress SCI-triggered neuroinflammation. These results indicated that sustained MSC-EXO delivery was the key to efficiently alleviating the neuroinflammatory microenvironment in spinal lesions after SCI^43^.

To further analyze the tissue-protective efficacy, sagittal sections of injured spinal cords at 8 weeks post-SCI were analyzed with immunofluorescence staining. As a result, the section images with Iba1 (macrophages/microglia) staining indicated that the activated immune cells in the rats were reduced significantly with MN-MSC patch treatment compared with those of the rats in the control group or that had the MN-EXO treatment (Supplementary Fig. S4a, e). In contrast, more neurons (NeuN) (Supplementary Fig. S4b, f), blood vessels (CD31) (Supplementary Fig. S4c, g), myelin (MBP) (Supplementary Fig. S4d, h) and astrocytes (GFAP) (Supplementary Fig. S4a-d, i) were observed in spinal tissues of the rats in the MN-MSC group. These results further verified that MN-MSC patch treatment could reduce SCI-triggered inflammation and protect neurons, oligodendrocytes, astrocytes and blood vessels from secondary injury after SCI.

### 3.4 The MN-MSC Patch Protects Spared Tissues/Axons from Secondary Injury

After verifying that MN-MSC treatment could reduce SCI-triggered inflammation, we sought to assess whether this treatment could protect spared tissues/axons from secondary injury after SCI. To determine whether more descending axons could survive after being treated with an MN-MSC patch, we injected AAV2/9-hSyn-mCherry into the T7-T8 region of the 3 groups of rats 2 weeks before sacrifice to trace the descending propriospinal axons (Fig. 3a). First, we found that the spinal cord shape deformation treated by the MN-MSC patch after 8 weeks was significantly less than that of rats in the control and MN-EXO groups (Fig. 3b). After staining the spinal sections with a red fluorescent protein (RFP) antibody (Fig. 3c), we found that the RFP^+^ propriospinal axons exhibited no significant difference in coronal sections of spinal segments above the injury site of rats in the three groups (Fig. 3e). However, the RFP+ propriospinal axons of rats in the MN-MSC group were higher than those of rats in the other groups, as indicated by the images of coronal sections of spinal segments below the injury sites (Fig. 3f).

**Figure 3.**
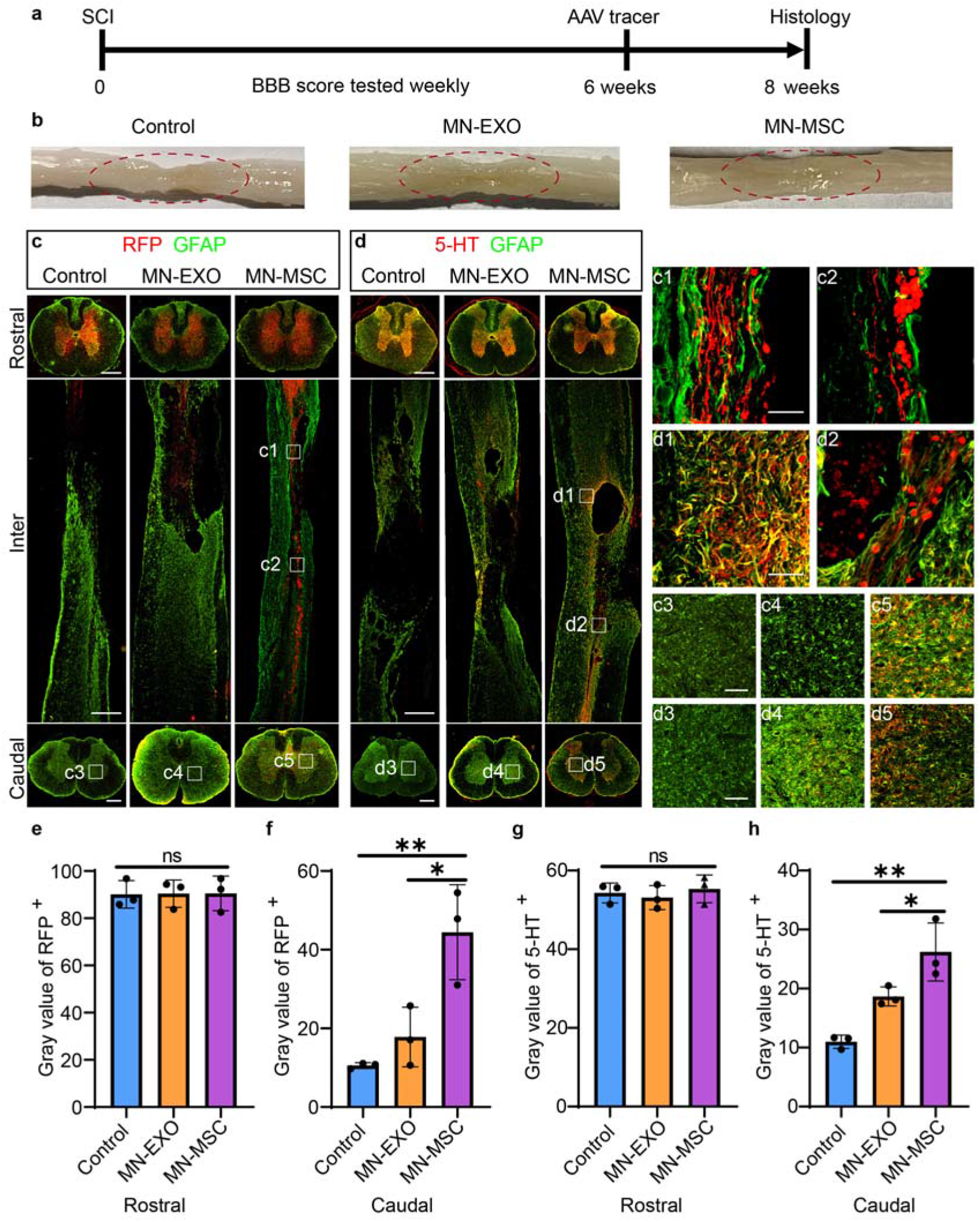
Protection of the spared axons with MN-MSC patch treatment after SCI. a) Schematic diagram of the experimental design. The MN or MN-MSC patch was implanted 3 hours after SCI, the AAV tracer for axons was injected at 6 weeks after injury, and the histological study was performed at 8 weeks after injury. The locomotion performance of the rats was recorded once a week during the experiments. b) Optical images of injured spinal cords of the three groups (control, MN-EXO, and MN-MSC) at 8 weeks after injury. Black arrows indicate the injury site. The regions of the injury site are marked with red dotted lines. c, d) Representative images of the spinal sections stained with GFAP (green), RFP (red), and 5-HT (red) of rats in the three groups (control, MN-EXO, and MN-MSC) at 8 weeks after SCI. Scale bars of rostral and caudal indicate 500 μm, Inter indicate 1 mm, c1, c2, d1, d2) indicate 100 μm, c3-c5, d3-d5) indicate 100 μm. e-h) Quantification of the gray values of 5-HT and RFP immunoreactivity on the rostral and caudal sides of the three groups. Data are shown as the mean ± SEM. Statistical analysis was performed using one-way ANOVA followed by Tukey’s multiple comparisons test. Statistical analysis was performed using one-way ANOVA followed by Tukey’s multiple comparisons test. n = 3. *p < 0.05, **p < 0.01.

Then, we stained the serotonergic axons and ubiquitous axonal fibers with anti-5-hydroxy tryptamine (5-HT) and anti-neurofilament (NF) in the spinal sections, respectively. We found that the trends of 5-HT and NF staining results were similar to that of RFP staining (Fig. 3d, S5a). No significant difference between NF and 5-HT was found above the injury site in the three groups (Fig. 3e, S5b). In contrast, the rats in the MN-MSC group had more serotonergic axons (stained by 5-HT) and axonal fibers (stained by NF) below the injury site than the rats in the other two groups (Fig. 3h, S5c), indicating that more propriospinal axons survived after SCI with MN-MSC patch treatment. In contrast to the control and MN-EXO patch-treated rats, we only found axons that extended into the spinal lesion site in the examined subjects with MN-MSC treatment (Fig. 3c, 3c1, 3c2, 3d, 3d1, 3d2, S5a, S5a1, S5a2), suggesting that the microenvironment with MN-MSC patch treatment was more conducive to axon regrowth. In addition, we did not observe apparent effects of the MN-MSC patch treatment on the other organs (Supplementary Fig. S6), suggesting that this treatment is safe for treating SCI.

### 3.5 The Remnants of Spared Axons Mediate Hindlimb Locomotor Functional Recovery

After verifying that more spared axons could be rescued by MN-MSC patch treatment, we sought to determine whether these axons could mediate hindlimb locomotor functions. To this end, we analyzed the hindlimb performance of the rats (three groups) with the Basso, Beattie, and Bresnahan (BBB) scale in a double-blind manner. We found that the rats in the control and MN-EXO groups exhibited paralysis with occasional ankle movement during the observation period, with an average BBB score of less than 4 (Fig. 4c). Even though the rats in the MN-MSC group predominantly demonstrated paralyzed hindlimbs during the first 4 weeks and showed no significant difference from the control and MN-EXO groups (Fig. 4c), they showed vigorous knee and ankle movement at 5 weeks. The BBB scores of these rats were significantly higher than those of rats in the other two groups 5 weeks postinjury. More strikingly, at the 8^th^ week after SCI, the rats treated with the MN-MSC patch showed improved hindlimb motor function, with higher body weight support and stride length than the rats in the other two groups (Fig. 4c, d, f, g). Remarkably, we found that 4 of 10 rats in the MN-MSC patch-treated group showed sustained weight support with hindpaw plantar stepping, and the BBB score reached more than 10, which was not observed in the rats in the control or MN-EXO groups (Fig. 4b, Supplementary Movie S1). Moreover, through subsequent analysis of the 7 kinds of hindlimb kinematics of rats in the control, MN-EXO, MN-MSC and intact groups, we found that the MN-MSC patch treatment could increase the maximal height of the iliac crest and toe (Supplementary Fig. S7a, b); the height amplitude of the iliac crest and toe (Supplementary Fig. 7c, d); and the angle oscillation of the hip, knee and ankle (Supplementary Fig. S7e-g). The hindlimb locomotor performance of rats in the MN-EXO group was slightly better than that of rats in the control group, but the difference was not significant (Supplementary Fig. S7a-d). However, the hindlimb locomotor performance of the rats treated with the MN-MSC patch was significantly better than that of rats in the MN-EXO and control groups but not as good as that of the rats in the intact group, suggesting that the hindlimb functional recovery with this treatment relied on the remnants of spinal connections across the lesion.

**Figure 4.**
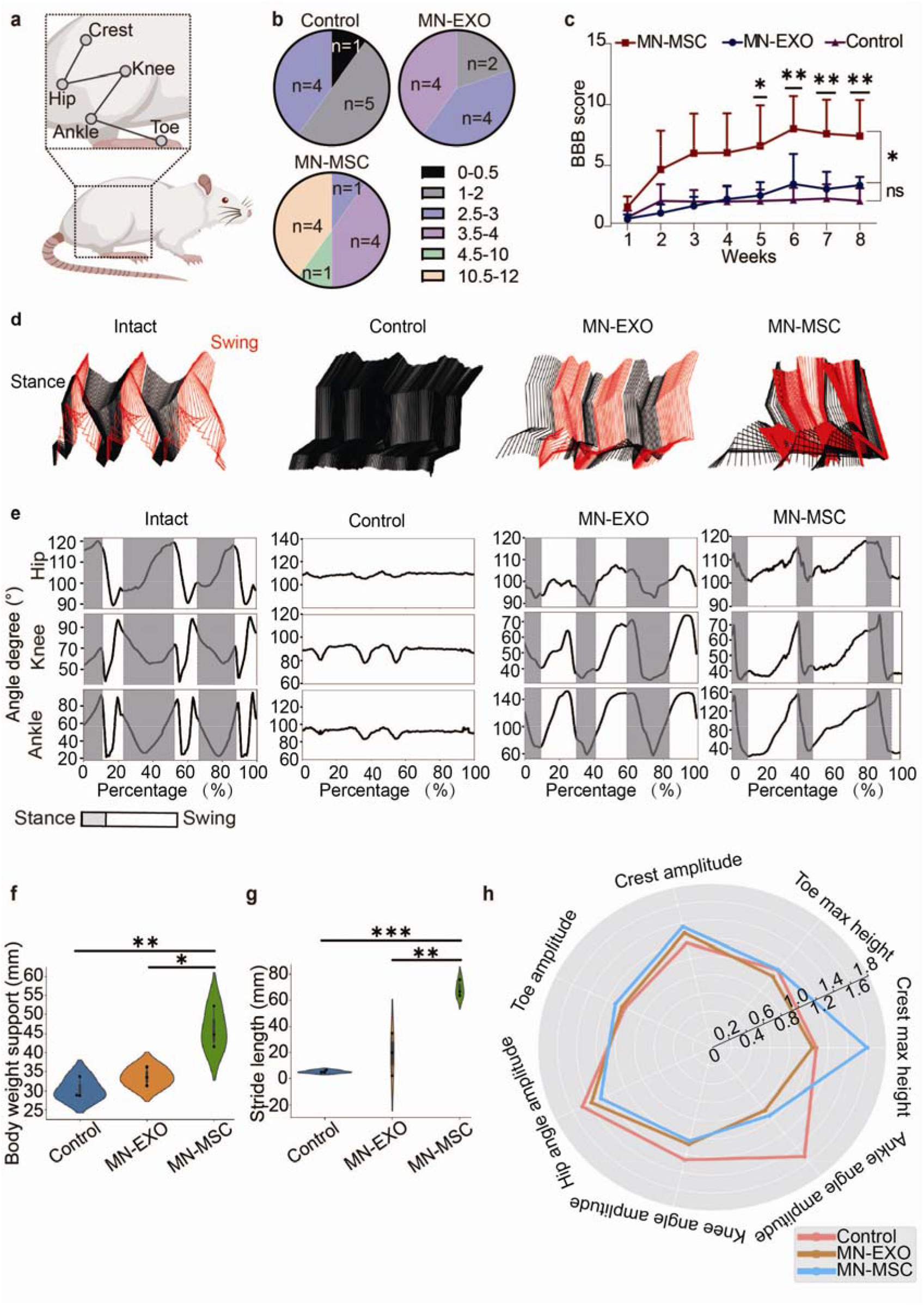
The hindlimb locomotor functional recovery of rats that underwent MN-MSC patch treatment after SCI. a) Schematic illustration of the position of hindlimb crest, knee, hip, ankle and toe. B) The distribution of BBB scores of rats in the three groups (control, MN-EXO, and MN-MSC) at 8 weeks after SCI. c) The weekly BBB score comparison among the rats in the three groups (control, MN-EXO, and MN-MSC) at 8 weeks after SCI. One-way ANOVA with Tukey’s post-hoc test was used for comparisons among multiple groups, and two-tailed paired t tests were used for comparisons between two groups. n=10. *p<0.05 and **p<0.01. d) The color-coded stick views of kinematic hindlimb analysis of the intact rats and the rats with SCI in the control, MN-EXO, and MN-MSC groups. e) The angle degree curves of the ankles, knees and hips of the intact rats and the rats with SCI in the control, MN-EXO, and MN-MSC groups. Quantitative analysis of f) body weight support and g) strike length at 8 weeks after injury. Data were analyzed with one-way ANOVA followed by Tukey’s post-hoc test. n = 3. *p < 0.05, **p < 0.01, ***p < 0.001. h) Well-rounded quantification of seven described behavioral features of rats in the three groups (control, MN-EXO, and MN-MSC), subjected to the indicated treatments through a radar graph.

To take both features of data distribution and statistical meaning into account in each kinematics indicator, we drew a radar plot (Fig. 4h). While all indicators in each group were divided by the counterpart in the intact group, it was obvious that the amplitudes of the crest and toe of the rats in the MN-MSC group were fairly high compared with those of rats in the MN-EXO and control groups (Fig. 4h). However, the maximum heights of the toe and crest of the rats treated with MN-MSC were remarkably higher than those of rats in the MN-EXO and control groups (Fig. 4h), possibly because the crest and toe determine the gait and stride length while walking. Comprehensively considering the value and distribution in each group, we found that the angle amplitudes among the ankles, knees and toes of rats in the MN-MSC group remained low, which is consistent with the results of the hindlimb kinematic study (Supplementary Fig. S7a-g).

To further determine the function of the MN-MSC patch treatment, we performed a somatosensory evoked potential (SSEP) monitoring study of the rats during anesthesia (Fig. 5a). Compared with the BBB score, SSEP is more direct for measuring the functionality of neuropathways from cortical projections^44^. As expected, we found that the SSEPs showed a similar trend to the BBB scores at the 8^th^ week postinjury (Fig. 5b, c), which confirms the axon protection capability of MN-MSC patch treatment. Then, we recorded electromyography (EMG) data to assess the rats’ hindlimb muscle motion when they were walking. We found that the ankle flexor signals of tibialis anterior (TA) muscles and extensor gastrocnemius soleus (GS) muscles were rarely active in rats in the control group, and the muscle signals of MN-EXO-treated rats could only be occasionally observed. In contrast, alternating activation of the TA and GS for the MN-MSC patch-treated rats could be consistently observed during different steps (Fig. 5d-f). To further analyze the EMG data, astronomical analysis (Poincaré statistical analysis) was used to analyze the rhythm of the TA and GS muscles. We found that the rhythms of TA and GS muscles of the rats in the MN-MSC group were higher than those of rats in the control and MN-EXO groups with a wider distribution range of scatter points and more concentrated at the central points (zero) position, which were more similar to the ones of the intact group (Supplementary Fig. S8), suggesting that those rats gain much better muscle control through MN-MSC patch treatment. From this, we explored why the gaits of rats treated with MN-MSC patches were closer to normal than the gaits of other rats. Together, these results suggested that rescued axons by MN-MSC patch treatment could mediate robust hindlimb locomotor functional recovery.

**Figure 5.**
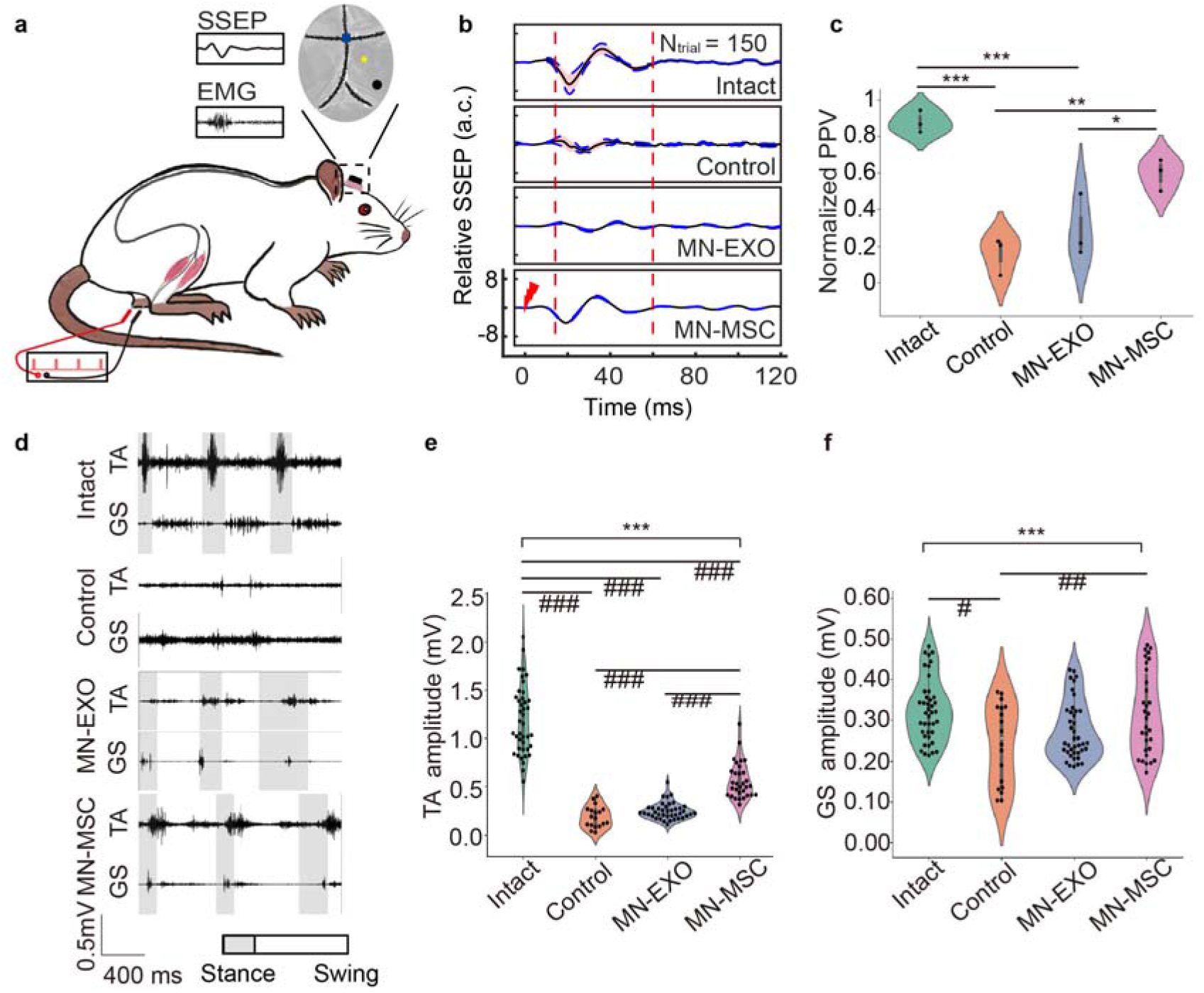
Electrophysiological recording experiments revealed the superior muscle control of rats treated with the MN-MSC patch. a) Leads were connected to a pair of antagonistic muscles (TA; medium GS) and the epidural electrode above the cortical hindlimb area (marked as a yellow star, B-2/R+2.5/D-0.7) for cortical evoked potential (stimulated by a pair of electrodes on the contralateral hindlimb paw) and free-moving muscle activity, respectively (blue triangular, bregma; black circular, ground lead). b) Representative relative SSEPs of each group (N trial = 150) are illustrated as the mean (solid black line) and 95% confidence interval (pink shadow enveloped by a blue dashed line), and the time of electric stimulation was set to zero ms. c) Peak-to-peak potential was analyzed with one-way ANOVA followed by Tukey’s post-hoc test. n = 3. *p<0.05, **p<0.01, ***p<0.001. d) TA and GS muscle EMG for rats of different groups (intact, control, MN-EXO and MN-MSC). The representative muscle activity of each group is depicted as three consecutive steps, where the gray intervals refer to stance phases. e, f) Quantitative analysis of signal amplitudes from TA and GS muscles for rats in different groups (intact, control, MN-EXO and MN-MSC). One-way ANOVA with Tukey’s post-hoc test for comparisons among multiple groups (*) and two-tailed paired t tests were used for comparisons within groups (#) for the data shown in the violin plot. n = 18-41. ***p (or ###p) < 0.001, **p (or ##p) < 0.01, *p (or #p) < 0.05.

## 4. Discussion

In this study, we constructed a novel MN-MSC patch that can provide sustained delivery of MSC-EXOs from the MSC-embedded patch to the spinal tissues in the injury site and avoid the risk of direct stem cells invading the spinal cord. The MN component of the patch was made of porous GelMA hydrogel with good biocompatibility, and the patch can remain on top of the spinal lesion for a long time, thus encompassing the optimal therapeutic window and achieving the maximum therapeutic efficacy of MSC-EXO treatment.

The study proposed a novel MN-MSC patch for SCI treatment, which could prevent additional surgical damage to the injured spinal cord and provide sustained highly effective MSC-EXO release. The retention time of MSC-EXOs in a living subject is short, but treatment with MSC-EXOs for at least 7 days after SCI is required to prevent secondary injury^44^. Moreover, the direct injection of MSC-EXOs or biomaterials containing MSC-EXOs into the lesion of an acute SCI is not ethically acceptable due to unpredictable clinical accidents. In comparison with these traditional treatment methods with MSCs or MSC-EXOs for SCI, the MN-MSC patch seeded with MSCs could avoid this ethical issue and provide an ethically acceptable method to achieve sustained delivery of MSC-EXOs for at least 7 days after SCI, which efficiently modulates dysregulated neuroinflammation after SCI and protects the spared tissue/axons from secondary injury^42^. In comparison with the traditional local injection methods, the method provided by this study topically delivers MSC-EXOs safely to the spinal injury site through the porous MN structure with higher accuracy and efficiency than other methods^45^.

Our findings showed that MN-MSC patch treatment could promote functional recovery by protecting the spared axons from secondary injury. In this study, the results of anatomical and functional recovery were analyzed in a double-blinded manner, ensuring the authenticity and effectiveness of the data. Consistent with the MSC-EXO function of immune modulation, apoptosis inhibition and enhanced angiogenesis^18,19,46^, more vessels (CD31, Arg-2) formed in the spinal lesion of SCI rats with our treatment; such vessels can supply sufficient nutrients and oxygen to remodel the SCI microenvironment and then protect spared axons^50,51^. Furthermore, the spinal cord of rats treated with the MN-MSC patch showed less inflammation (Iba1, MMP-9) with more neurons (NeuN), less apoptosis and death of spinal cord cells (Bax), and more myelin sheath (MBP), which benefit the plasticity of spinal circuitry in terms of function. By axon tracing and labeling, we showed that more axons survived secondary injury by MN-MSC patch treatment. In comparison with the rats in the control or MN-EXO treatment groups, the rats with MN-MSC patch treatment exhibited remarkable hindlimb locomotion functional recovery, which is consistent with their better muscle control ability, as indicated by the PPV and signal of TA and GS muscle EMG recording studies. However, a variety of functional recovery levels was observed in the rats treated by MN-MSC patches, possibly because the rescued axons that are closely relevant to hindlimb locomotion were randomly presented in the treated rats.

Even though we demonstrated the proof-of-concept of the MN-MSC patch for SCI treatment, the optimal time window for MN-MSC patch implantation remains to be examined. Since the MN-MSC patch was fabricated from GelMA hydrogel, we envisioned that the MN-MSC patch would degrade in the long term^47,48^. However, whether and when the materials of the MN-MSC patch could completely degrade should be investigated in future studies. In addition, the relationship between the number of exosomes released from the MN-MSC patch and the best treatment efficacy has not been tested and therefore needs to be investigated before clinical study. To promote diffusion and improve delivery efficiency, in our next step, we plan to construct a precise porous MN structure at the micro/nano scale with two-photon 3D printing technology, which might have higher precision in controlling the porous size of the MNs^49^. Moreover, the MN-MSC patch might be too soft to be used as an artificial dura, and a tenacious MN-MSC patch with the addition of a scaffold structure between the MN array and the MSC-embedded patch is being developed by our group.

## Methods

### Materials

Chicken anti-GFAP [abcam (ab1344360, 1:500)], rabbit anti-neurofilament (NF) heavy polypeptide [abcam (ab8135), 1:500], goat anti 5-HT antibody [Invitrogen (pa1-36157), 1:500], rabbit anti-NeuN [abcam (ab177487), 1:1000], rabbit anti-MBP [abcam (ab218011), 1:500], goat anti-IBA1 [abcam(ab5076), 1:500], rabbit anti-CD31 [R&Drd system (AF3628), 1:500], rabbit anti hHuman Specific GAPDH Rabbit mAb [ABclonal (AB 2769571,1:500)], rabbit anti-TGF-β [Abclonal(A15103), 1:1000], rabbit anti-Bax [Abclonal (A0207),1:1000], rabbit anti-MMP9 [Abclonal (A0289), 1:1000], rabbit anti-Arginase 2 (ARG2) [Abclonal (A19233), 1:1000], rabbit anti-GAPDH [Abclonal (AC001), 1:1000], Donkey anti-Chicken IgY H&L (FITC) [Abcam (ab63507), 1:500], Donkey anti-Rabbit IgG H&L (Alexa Fluor^®^ 555) [Abcam (ab150062), 1:500], Rabbit anti-Goat IgG H&L (Alexa Fluor^®^ 555) [Abcam (ab150142), 1:500] and Donkey anti-Rabbit secondary antibodies (HRP) [beyotime (A0208), 1:1000] were used. The adenovirus-associated virus AAV2/9-hSyn-mCherry was generated by the viral core of Zhejiang University, and its titer was adjusted to 1 × 10^13^ copies per mL for injection. DAPI-containing antifade mounting medium was purchased from Southern Biotech (0100-20, USA). Triton X-100 (P1080) was purchased from Solarbio (China).

### Animals

Female Sprague□Dawley rats (220-250 g) were purchased from the Zhejiang Academy of Medical Sciences. All experiments were approved by the Zhejiang University School of Medicine Animal Experimentation Committee (approval ID: ZJU20210110). The rats were housed under controlled environmental conditions and were completely in compliance with the National Institutes of Health Guide for the Care and Use of Laboratory Animals. All surgeries were performed under anesthesia with 1% (w/v) pentobarbital sodium (5 mL/kg, injected intraperitoneally).

### Preparation of MSC

The human embryonic stem cell (ESC)-derived MSCs were gifts from Ysbiotech, Hangzhou, China (YS^™^ hESC-MSC). The MSCs were generated from an H9 human embryonic stem cell line and cultured at 37°C in a humidified atmosphere of 5% CO_2_ in serum-free medium for MSCs (NC0103+NC0103. S, Yocon, Beijing, China) as previously reported^50^, with some modification. The H9 human embryonic stem cell line, originally generated by National Stem Cell Bank c/o WiCell Research Institute (USA), was purchased from the stem cell bank at Institute of Biochemistry and Cell Biology, CAS. When the proliferating colonies had reached near confluence, the hESC-MSCs were passaged using stem cells moderate digestive enzymes (NC1004, Yocon, Beijing, China). After 3-4 passages, MSCs were used for transplantation or other investigations.

### Fabrication and Characterization of MN Arrays

The MN arrays were fabricated with polydimethylsiloxane molds (EFL-MMN-600) from EFL Ltd. The MNs had diameters of 250 μm, which tapers for a height of 600 mm. The MN arrays were arranged 550 μm, tip to tip, in a 4 mm×4 mm area. First, 5% porous GelMA (EFL-GM-pr-001) solution was prepared in a 37°C water bath and dropped into the molds. Next, the bubbles of the casting solution were removed by vacuum pumping, and the molds were placed in a 37°C environment for 10 hours to dry. Then, the casting solution was solidified with blue light curing (405 nm, 30 mW/cm2) for 30 s. After that, the MN was separated from the molds with tweezers cautiously. The MN array was cut into small pieces at a scale of 3 mm×3 mm. The morphology of the MNs was assessed with a SEM (Nova Nano 450, Thero FEI, USA).

### Fabrication and Characterization of the MN-MSC Patch

First, the MSCs were digested by stem cell digestive enzymes (NC1004.1) and evenly mixed in 5% porous GelMA solution at a ratio of 2.5×107 cells/mL at 37°C. Next, the solution was added to the basal side of the MN arrays, and the MN-MSC patch was obtained with a blue light curing process for 30 s. The MN-MSC patch was cultured in MSC serum-free media.

The viability of MSCs in the MN-MSC patch was evaluated with a live/dead viability/cytotoxicity kit (L3224, Invitrogen, USA). After washing the MN-MSC patch with phosphate buffered saline (PBS) for three times, the MSCs were stained with 0.2% ethidium homodimer-1 and 0.05% calcein AM in PBS for 20 min. Then, the MN-MSC patch was washed three times and observed by laser scanning confocal microscopy (Nikon, A1R). The proliferation of the MSCs in the MN-MSC patch was tested by CCK-8 assay (CK04, Dojindo, Japan) on the 1^st^, 3^rd^ and 5^th^ days. And the optical density of formazan was detected at a wavelength of 450 nm by a microplate reader (iD5, Molecular Devices, USA). MN-EXOs were fabricated in a similar way to MSC-secreted exosomes (1.25 μg/μl) instead of MSCs.

### Surgical Procedure

For the *in vivo* experiments, a spinal contusion injury model was induced by an infinite vertical impactor (68099, RWD, China). Specifically, after being anesthetized, a laminectomy was performed at the tenth thoracic vertebral level (T10-11) of a rat to expose the dorsal surface to induce SCI^51^. Next, contusion was performed using a cylinder (diameter, 3 mm) that impacted the spinal cord at a rate of 2.5 m/s and was left for 5 s at a depth of 2 mm after being impacted (Supplementary Fig. S2a, b). After 3 hours, the dura mater at the lesion site was cut carefully using microscissors (S11001-09, RWD, China) in the direction of the spinal cord, and an MN was placed at the site of the dura matter defect (Supplementary Fig. S2c, d). GelMA with MSCs (25,000 cells/μL) was dripped onto the MN and irradiated with light at a wavelength of 405 nm for 1 minute to create the MN-MSC patch (Supplementary Fig. S2e). Then, more 5% GelMA solution was dripped on the surface of the exposed spinal cord with the patch and gelled with blue light to fix the patch on the injury site of the spinal cord (Supplementary Fig. S2f-h). The control group rats underwent contusion and dura mater cutting, but no MN was placed. The MN-EXO implantation surgery was performed in the same way with MSC-secreted exosomes (1.25 μg/μl) instead of MSCs. Artificial micturition was provided twice daily until automatic bladder voiding was restored.

At 6 weeks after injury, anterograde tracing of propriospinal axons was performed via the injection of AAV2/9-hSyn-mCherry at the T7-8 spinal cord^42^. Specifically, a pull-glass micropipette tipped with a 10-μL Hamilton microsyringe (68606, RWD, China) was used for precise injection. After a laminectomy was performed at the T7-8 spinal cord, AAV2/9-hSyn-mCherry was injected into 12 sites at the following locations: 1) 0.4 and 0.8 mm lateral to the midline; 2) 0.5, 1.0 and 1.6 mm from the surface. The injection rate was 80 nl/min, and the injected liquid volume was 150 nl per site. The needle was left for 1 minute before moving to the next. Animals were sacrificed for assessment 2 weeks after injection.

### Evaluation of the MSC-EXO Release Profile of the MN-MSC Patch

To evaluate the advantage of the release function of the MN-MSC patch, we measured the MN-EXO patch release capacity (containing EXOs instead of MSCs) against the capacity of our MN-MSC patch. The patches were cultured and placed on the surface of culture inserts (Fig. 1i) to investigate the MSC-EXO release function of the two different patches. The MSC-EXOs released in the culture medium were quantified with a Micro BCA Protein Assay kit (P0012, Beyotime, China).^39^ The MSC-EXOs were collected from the supernatant using ultracentrifugation. The size and morphology of MSC-EXOs were tested with transmission electron microscopy (TEM) (JEM-1400flash JEOL, JEOL, Japan).

Because the MSC-EXOs released from the MN-MSC patch were difficult to examine visually, we used the CM-DiI-MN-EXO patch [EXOs labeled with CM-DiI (2 mg·mL-1, Yeasen, China)] to observe the *in vivo* release capability of the MN patch. The CM-DiI-MN-EXO patch was implanted in the injured spinal cord of the rat. The method of patch implantation was described in the animal and surgical procedures section. The stained images of the EXOs released in the SCI were acquired by a slide scanner (VS200, Olympus, Japan).

### Histochemistry

Animals were sacrificed at 8 weeks after injection for histological assessment. After the animals were anesthetized, perfusion was performed using PBS and 4% paraformaldehyde. Spinal cords and other tissues were fixed in 4% paraformaldehyde overnight and dehydrated with sucrose for 1 day. After being embedded using optimal cutting temperature compound (OCT), twenty-micrometer-thick sections of the spinal cord were cut using a cryostat (CryoStar NX50; Thermo, USA) and mounted onto slides.

For immunofluorescence histochemistry, spinal cord tissue sections were blocked with 5% donkey serum and 0.3% Triton X-100 for 1 hour and incubated with primary antibodies overnight at 4°C. After washing with PBS three times, the tissues were incubated with secondary antibodies conjugated to fluorescent dyes for 2 hours at room temperature. Then, tissues were rewashed with PBS and dripped with an antifade mounting medium containing DAPI. Images were acquired by a confocal laser scanning microscope (A1Ti, Nikon, Japan) and a slide scanner (VS200, Olympus, Japan).

For immunohistochemistry, tissue sections were stained with H&E. Images were acquired by a slide scanner (VS200, Olympus, Japan).

### Western Blotting

Cells were washed three times with ice-cold PBS and lysed in RIPA buffer containing 1% PMSF. After leaving the samples on ice for 30 minutes, they were centrifuged at 12000 × g for 15 minutes at 4°C. Protein concentrations in the supernatant were assessed using a Micro BCA Protein Assay kit. Equal amounts of protein extracts were resolved by 10%-12% SDS-PAGE and electrotransferred onto a polyvinylidene membrane (Bio-Rad, USA). After blocking in Tris-buffered saline plus 5% (w/v) milk, the membranes were exposed to primary antibodies overnight at 4°C. The samples were incubated with secondary antibodies conjugated to horseradish peroxidase for 1 hour at room temperature. Signals were visualized by a ChemiDoc Touch Imaging System (Bio-Rad, USA), and ImageJ software was used to assess gray levels. All experiments were repeated three times.

### Behavioral Assessment

The locomotion performance of the rats was recorded by a camera once a week during the experiments. An examiner blinded to the treatment assessed the videos recording the rats’ behavioral performance by the Basso, Beattie Bresnahan rating scale (BBB scale)^52^. Rats that showed a BBB score above 1.5 at 1 week after SCI were excluded from further histological or behavioral analysis. To precisely understand the locomotor kinematics, the representative hindlimb osteoarticular landmarks (toe, ankle, knee, hip, and crest) were labeled by 5 mm reflective balls and tracked by the MotoRater (Vicon Nexus, UK)^42^. Typical sequential treads and the ankle degrees of selected articulates were displayed. Body weight support and stride length were compared as previously described^52^.

The behavioral data were depicted with seven features (maximal iliac crest height; crest height amplitude; maximal toe height; toe height amplitude; and hip, knee and ankle angle oscillation) within four groups (intact, control, MN-EXO, and MN-MSC). For each group, we utilized different statistical methods to evaluate the seven behavioral features, and these features have different units. As we needed to thoroughly analyze these features in radar graphs, normalizing these data was vital and pivotal. First and foremost, for each value in each feature, we normalized the behavioral data of the seven features by equation (1):

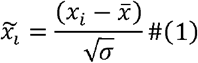

where *x_i_* represents the *ith* data point for each of the seven behavioral features, *i* = 1,2,3,…*n*. *n* is the number of points sampled from each feature in each group. 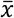 and 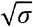 represent the mean value and standard variance for each feature, respectively.

Second, we obtained the seven 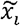 values. Each group had positive and negative values, and the scale among these values was not fixed. Hence, we projected the features into an exponentiation expression, mapping these values in a unified scale (a unified vector space). For each feature depicted in a different group, we used equation (2) to process this step:

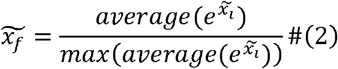

where *f* = 1,2,3,…,7, represents the seven behavioral features in each group. We averaged the 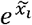 in each group, obtaining lists with seven feature values in each group.

In the final analysis, we used the intact group values as the benchmark, significantly depicting the feature values’ variance. We used equation (3) to process this step:

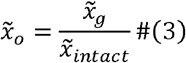

where 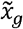 represents each feature value in each group. 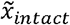 represents the feature values in the intact group, namely, the benchmark values.

### Implantation of Electrodes

Electrode implantation was performed 8 weeks after SCI for both evoked potential and hindlimb EMG according to previously published protocols^42^. Rats were anesthetized by dexmedetomidine hydrochloride (0.25 mg/kg i.p.; MCE, US) and 0.5% isoflurane, which worked well on the acquisition of evoked potentials^53^. A stainless steel screw electrode was fixed on the skull above the hindlimb region stereotactically (B-2.0/R+2.5/D-0.7) for an epidural recording of cortical evoked potentials referring to the zero potential on the shoulder (ground lead connected to B-6.0/R4.0)^54^. Following the SSEP recording, bipolar electrodes (AS632, Conner wire) were inserted into the right medium gastrocnemius (GS) and TA for *in vivo* assessment of hindlimb muscles with the reference lead linked to the ipsilateral tendon^42^. Wires were subcutaneously connected to the header plate fixed on the skull.

### SSEP Recording and Processing

During the surgery, the header was connected to a neuron signal amplifier (BTAM01L, Braintech, China), and the signal was recorded by a NeuroStudio system (Braintech, China)^42^. A series of pulses (2.0 Hz, 2.0 mA, 0.5 ms) was applied to the left hindlimb paw through a pair of electrodes by a neurostimulator (BTSEM-16, Braintech, China) under anesthesia^53^. Signals were obtained from a five-minute stimulation, followed by a 30-to 1500-Hz bandpass, power line noise notch filtration (50 Hz and the two consequent harmonics 100 Hz and 150 Hz), and the removal of stimulation artifacts. Then, the first 150 qualified trials were chosen for peak-to-peak potential (PPV) calculation. The PPV was normalized according to the intact group, following the method described by Gunnar Waterstraat et al^55^.

### EMG Recording and Processing

Seven days postimplantation, five osteoarticular structures were captured to distinguish stance and swing when moving freely, and the header was connected up to the amplifier to detect muscle strength. Next, the free-moving muscle activity, represented by an extensor and a flexor of the right hindlimb, was logged in NeuroStudio. Data were filtered (bandpass 20 Hz-1000 Hz) before computing the muscle amplitude on the MATLAB platform^51^.

### Statistics

Multiple samples were analyzed by one-way or two-way analysis of variance (ANOVA). Differences between the vehicle control and experimental groups were analyzed by Student’s t test. All parameters were expressed as the mean ± standard error of the mean (SEM), and a P value less than 0.05 was considered significant.

## Acknowledgments

This study was supported by the Scientific and Technological Innovation 2030 Program of China - major projects (2021ZD0200408 to X.W.), the National Natural Science Foundation of China (81971866 to X.W.), the Science Fund for Distinguished Young Scholars of Zhejiang Province (LR20H090002 to X.W.), the Leading Innovative and Entrepreneur Team Introduction Program of Zhejiang (2019R01007 to X.W.), the Fundamental Research Funds for the Central Universities (K20210195 to X.W.) and the Postdoctoral General Foundation project (2020M671746 to X.C.).

## Author Contributions

A.F., Y.W., N.G. contributed equally to this work. X.W. and A.F. conceptualized and designed the study. A.F., Y.W., N.G. conducted the experiments and collected the data. A.F., Y.W., and N.G., L. L, B. G., W. C., X.C., J. Y., Z. A., X. L., Y. Z., H. Z., Z. W., S. J., K. X., X. G., B.Y. and X.W. analyzed and interpreted the data. A.F. and X.W. drafted the paper. All authors critically revised the manuscript and approved the final version for submission.

## Data Availability Statement

The data to support the findings of this study are included in the paper and supplementary information, and further data are available from the corresponding author upon reasonable request.

---

**Scheme 1.**
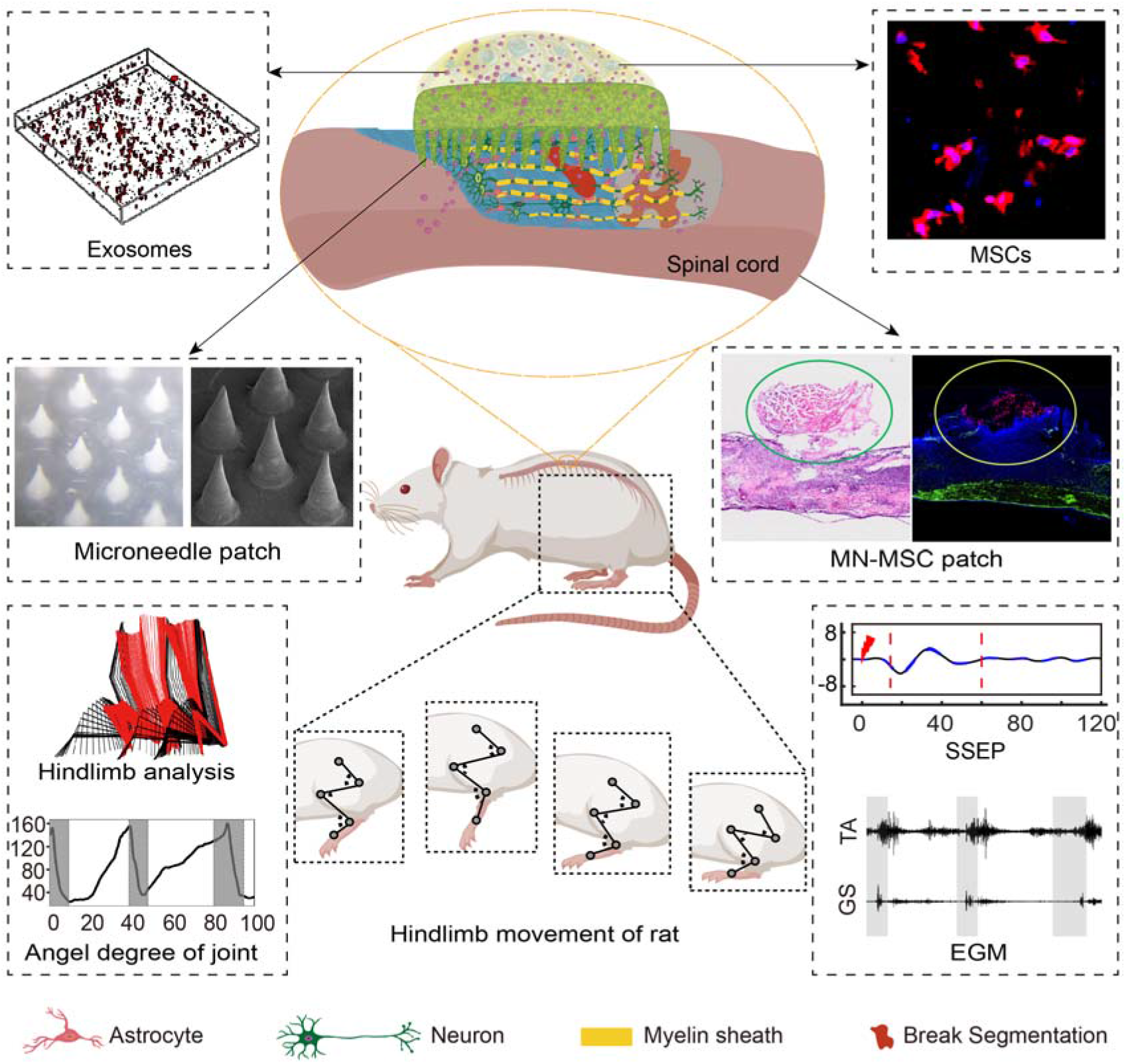
Schematic illustration of MN-MSC patch implantation at the site of the spinal cord injury.

## References

1 Collaborators, G. B. D. M. S. Global, regional, and national burden of multiple sclerosis 1990-2016: a systematic analysis for the Global Burden of Disease Study 2016. The Lancet. Neurology 18, 269–285, doi:10.1016/S1474-4422(18)30443-5 (2019).

2 Short, D. Use of steroids for acute spinal cord injury must be reassessed. Bmj 321, 1224 (2000).

3 Wu, Y. T. et al. Relationship between the interval before high-dose methylprednisolone administration and chronic pain in traumatic spinal cord injury. Neurosciences (Riyadh) 16, 324–328 (2011).

4 Matsumoto, T. et al. Early complications of high-dose methylprednisolone sodium succinate treatment in the follow-up of acute cervical spinal cord injury. Spine 26, 426–430, doi:10.1097/00007632-200102150-00020 (2001).

5 Christopher, S. et al. Traumatic spinal cord injury. Nature reviews. Disease primers (2017).

6 Bains, M. & Hall, E. D. Antioxidant therapies in traumatic brain and spinal cord injury. Biochimica Et Biophysica Acta 1822, 675–684 (2012).

7 Akhtar, A. M., Al, S. & Eid, A. H. Inflammogenesis of Secondary Spinal Cord Injury. Frontiers in Cellular Neuroscience 10 (2016).

8 Tahmasebinia, F. & Pourgholaminejad, A. The role of Th17 cells in auto-inflammatory neurological disorders. Progress in neuro-psychopharmacology & biological psychiatry, 408–416 (2017).

9 Shao, A., Tu, S., Lu, J. & Zhang, J. Crosstalk between stem cell and spinal cord injury: pathophysiology and treatment strategies. Stem Cell Research Therapy 10 (2019).

10 Huang, L., Fu, C., Xiong, F., He, C. & Wei, Q. Stem Cell Therapy for Spinal Cord Injury. Cell transplantation 30, 963689721989266, doi:10.1177/0963689721989266 (2021).

11 Yao, S. et al. Mesenchymal Stem Cell-Laden Hydrogel Microfibers for Promoting Nerve Fiber Regeneration in Long-Distance Spinal Cord Transection Injury. ACS Biomater Sci Eng 6, 1165–1175, doi:10.1021/acsbiomaterials.9b01557 (2020).

12 Wong, S. W., Lenzini, S., Cooper, M. H., Mooney, D. J. & Shin, J. W. Soft extracellular matrix enhances inflammatory activation of mesenchymal stromal cells to induce monocyte production and trafficking. Sci Adv 6, eaaw0158, doi:10.1126/sciadv.aaw0158 (2020).

13 Zhang, L. et al. NSCs Migration Promoted and Drug Delivered Exosomes-Collagen Scaffold via a Bio-Specific Peptide for One-Step Spinal Cord Injury Repair. Advanced healthcare materials 10, e2001896, doi:10.1002/adhm.202001896 (2021).

14 Nakazaki, M., Morita, T., Lankford, K. L., Askenase, P. W. & Kocsis, J. D. Small extracellular vesicles released by infused mesenchymal stromal cells target M2 macrophages and promote TGF-β upregulation, microvascular stabilization and functional recovery in a rodent model of severe spinal cord injury. Journal of extracellular vesicles 10, e12137 (2021).

15 Chhabra, H. S. & Sarda, K. Clinical translation of stem cell based interventions for spinal cord injury—Are we there yet? Advanced Drug Delivery Reviews 120, 41–49 (2017).

16 Ren, G. et al. Mesenchymal stem cell-mediated immunosuppression occurs via concerted action of chemokines and nitric oxide. Cell Stem Cell 2, 141–150 (2008).

17 Gnecchi, M. et al. Paracrine action accounts for marked protection of ischemic heart by Akt-modified mesenchymal stem cells. Nat Med 11, 367–368 (2005).

18 Zhang, S. et al. MSC exosomes mediate cartilage repair by enhancing proliferation, attenuating apoptosis and modulating immune reactivity. Biomaterials 156, 16–27 (2018).

19 Bian, S. et al. Extracellular vesicles derived from human bone marrow mesenchymal stem cells promote angiogenesis in a rat myocardial infarction model. Journal of molecular medicine 92, 387–397 (2014).

20 Rehman, J. et al. Secretion of angiogenic and antiapoptotic factors by human adipose stromal cells. Circulation 109, 1292–1298 (2004).

21 Li, L. et al. Transplantation of human mesenchymal stem-cell-derived exosomes immobilized in an adhesive hydrogel for effective treatment of spinal cord injury. Nano letters 20, 4298–4305 (2020).

22 Mu, J. et al. Hypoxia-stimulated mesenchymal stem cell-derived exosomes loaded by adhesive hydrogel for effective angiogenic treatment of spinal cord injury. Biomaterials Science 10, 1803–1811 (2022).

23 Ousman, S. S. & Kubes, P. Immune surveillance in the central nervous system. Nature neuroscience 15, 1096–1101, doi:10.1038/nn.3161 (2012).

24 Block, M. L., Zecca, L. & Hong, J. S. Microglia-mediated neurotoxicity: uncovering the molecular mechanisms. Nature reviews. Neuroscience 8, 57–69, doi:10.1038/nrn2038 (2007).

25 Lv, Z., Dong, C., Zhang, T. & Zhang, S. Hydrogels in Spinal Cord Injury Repair: A Review. Frontiers in bioengineering and biotechnology 10, 931800, doi:10.3389/fbioe.2022.931800 (2022).

26 Chen, C., Qiao, X., Liu, W., Fekete, C. & Reinhardt, J. D. Epidemiology of spinal cord injury in China: A systematic review of the chinese and english literature. Spinal cord, doi:10.1038/s41393-022-00826-6 (2022).

27 Li, J. & Mooney, D. J. Designing hydrogels for controlled drug delivery. Nat Rev Mater 1, doi:10.1038/natrevmats.2016.71 (2016).

28 Mukherjee, N., Adak, A. & Ghosh, S. Recent trends in the development of peptide and protein-based hydrogel therapeutics for the healing of CNS injury. Soft matter, doi:10.1039/d0sm00885k (2020).

29 Afferi, L. et al. Performance and safety of treatment options for erectile dysfunction in patients with spinal cord injury: A review of the literature. Andrology 8, 1660–1673, doi:10.1111/andr.12878 (2020).

30 Santi, S., Corridori, I., Pugno, N. M., Motta, A. & Migliaresi, C. Injectable Scaffold-Systems for the Regeneration of Spinal Cord: Advances of the Past Decade. ACS Biomater Sci Eng 7, 983–999, doi:10.1021/acsbiomaterials.0c01779 (2021).

31 Brommer, B. et al. Improving hindlimb locomotor function by Non-invasive AAV-mediated manipulations of propriospinal neurons in mice with complete spinal cord injury. Nature Communications 12, 781, doi:10.1038/s41467-021-20980-4 (2021).

32 Macaya, D. & Spector, M. Injectable hydrogel materials for spinal cord regeneration: a review. Biomedical materials 7, 012001 (2012).

33 Tang, J. et al. Cardiac cell–integrated microneedle patch for treating myocardial infarction. Science Advances 4, eaat9365 (2018).

34 Yang, G. et al. A therapeutic microneedle patch made from hair-derived keratin for promoting hair regrowth. Acs Nano 13, 4354–4360 (2019).

35 Zhou, X. et al. Biodegradable β-Cyclodextrin Conjugated Gelatin Methacryloyl Microneedle for Delivery of Water-Insoluble Drug. Advanced healthcare materials 9, 2000527 (2020).

36 Arya, J. et al. Tolerability, usability and acceptability of dissolving microneedle patch administration in human subjects. Biomaterials 128, 1–7, doi:10.1016/j.biomaterials.2017.02.040 (2017).

37 Dvorak, M. F. et al. Minimizing errors in acute traumatic spinal cord injury trials by acknowledging the heterogeneity of spinal cord anatomy and injury severity: an observational Canadian cohort analysis. J Neurotrauma 31, 1540–1547, doi:10.1089/neu.2013.3278 (2014).

38 Ying, G. et al. Bioprinted injectable hierarchically porous gelatin methacryloyl hydrogel constructs with shape-memory properties. Advanced functional materials 30, 2003740 (2020).

39 Li, L. et al. Transplantation of Human Mesenchymal Stem-Cell-Derived Exosomes Immobilized in an Adhesive Hydrogel for Effective Treatment of Spinal Cord Injury. Nano letters 20, 4298–4305, doi:10.1021/acs.nanolett.0c00929 (2020).

40 Branscome, H. et al. Use of Stem Cell Extracellular Vesicles as a “Holistic” Approach to CNS Repair. Front Cell Dev Biol 8, 455, doi:10.3389/fcell.2020.00455 (2020).

41 Brennan, M. Á., Layrolle, P. & Mooney, D. J. Biomaterials functionalized with MSC secreted extracellular vesicles and soluble factors for tissue regeneration. Advanced functional materials 30, 1909125 (2020).

42 Ye, J. et al. Rationally Designed, Self-Assembling, Multifunctional Hydrogel Depot Repairs Severe Spinal Cord Injury. Adv Healthc Mater 10, e2100242, doi:10.1002/adhm.202100242 (2021).

43 Ma, S. et al. Immunobiology of mesenchymal stem cells. Cell Death & Differentiation 21, 216–225 (2014).

44 All, A. H., Al Nashash, H., Mir, H. & Luo, S. Characterization of transection spinal cord injuries by monitoring somatosensory evoked potentials and motor behavior. Brain Research Bulletin 156, 150–163 (2020).

45 Qin, Y., Sun, R., Wu, C., Wang, L. & Zhang, C. Exosome: A Novel Approach to Stimulate Bone Regeneration through Regulation of Osteogenesis and Angiogenesis. International Journal of Molecular Sciences 17, 712 (2016).

46 Bruno, S. et al. Microvesicles derived from mesenchymal stem cells enhance survival in a lethal model of acute kidney injury. PloS one 7, e33115 (2012).

47 Heltmann-Meyer, S. et al. Gelatin methacryloyl is a slow degrading material allowing vascularization and long-term usein vivo. Biomed Mater 16, doi:10.1088/1748-605X/ac1e9d (2021).

48 Yue, K. et al. Synthesis, properties, and biomedical applications of gelatin methacryloyl (GelMA) hydrogels. Biomaterials 73, 254–271, doi:10.1016/j.biomaterials.2015.08.045 (2015).

49 Xing, J.-F., Zheng, M.-L. & Duan, X.-M. Two-photon polymerization microfabrication of hydrogels: an advanced 3D printing technology for tissue engineering and drug delivery. Chemical Society Reviews 44, 5031–5039 (2015).

50 Nakajima, H. et al. Transplantation of mesenchymal stem cells promotes an alternative pathway of macrophage activation and functional recovery after spinal cord injury. J Neurotrauma 29, 1614–1625, doi:10.1089/neu.2011.2109 (2012).

51 Chen, B. et al. Reactivation of Dormant Relay Pathways in Injured Spinal Cord by KCC2 Manipulations. Cell 174, 521–535 e513, doi:10.1016/j.cell.2018.06.005 (2018).

52 Basso, D. M., Beattie, M. S. & Bresnahan, J. C. Graded histological and locomotor outcomes after spinal cord contusion using the NYU weight-drop device versus transection. Exp Neurol 139, 244–256, doi:10.1006/exnr.1996.0098 (1996).

53 Liu, S. et al. Fully Passive Flexible Wireless Neural Recorder for the Acquisition of Neuropotentials from a Rat Model. ACS Sens 4, 3175–3185, doi:10.1021/acssensors.9b01491 (2019).

54 Iyer, S., Maybhate, A., Presacco, A. & All, A. H. Multi-limb acquisition of motor evoked potentials and its application in spinal cord injury. J Neurosci Methods 193, 210–216, doi:10.1016/j.jneumeth.2010.08.017 (2010).

55 Waterstraat, G., Körber, R., Storm, J. H. & Curio, G. Noninvasive neuromagnetic single-trial analysis of human neocortical population spikes. Proc Natl Acad Sci U S A 118, doi:10.1073/pnas.2017401118 (2021).

