## Supplementary figures for "Porous microneedle patch with sustained exosome delivery repairs severe spinal cord injury"

1. Department of Neurobiology and Department of Rehabilitation Medicine, First Affiliated Hospital, College of Medicine, Zhejiang University, Hangzhou, Zhejiang Province, 310003, PR China
2. NHC and CAMS Key Laboratory of Medical Neurobiology, MOE Frontier Science Center for Brain Research and Brain–Machine Integration, School of Brain Science and Brain Medicine, Zhejiang University, Hangzhou, Zhejiang Province, 310003, PR China
3. Department of Neurobiology and Department of Orthopedics, 2nd Affiliated Hospital, Zhejiang University School of Medicine, Zhejiang University, Hangzhou, Zhejiang Province, 310003, PR China
4. Key Laboratory of Neuroregeneration of Jiangsu and Ministry of Education, NMPA Key Laboratory for Research and Evaluation of Tissue Engineering Technology Products, Nantong University, Nantong, China
5. Co-innovation Center of Neuroregeneration, Nantong University, Nantong, 226001 Jiangsu, PR China
6. These authors contributed equally

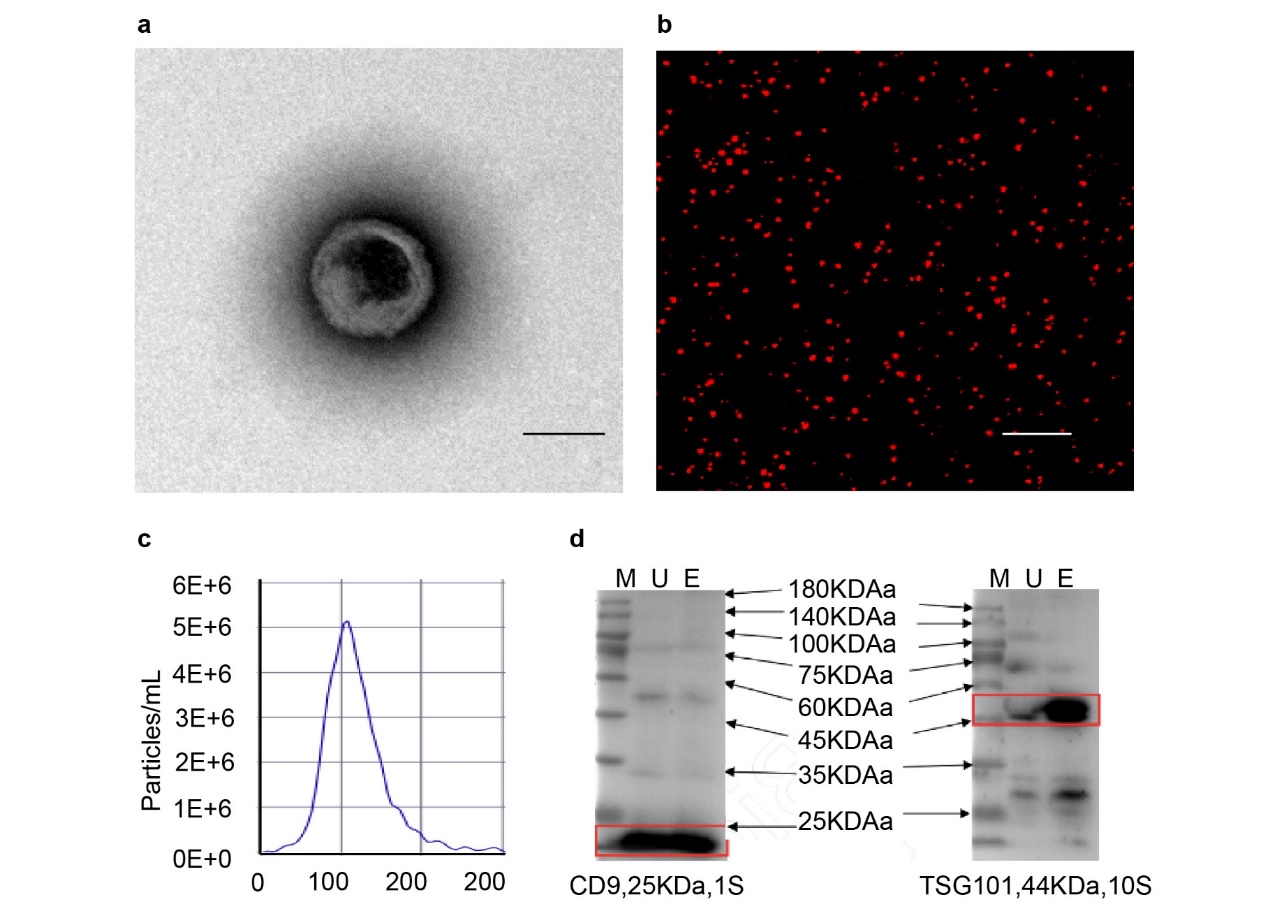

**Figure S1**. Characterization of exosomes excreted by MSCs. a) Morphology of exosomes revealed by TEM. Scale bar: 100 nm. b) Exosomes labeled by CM-DiI (red) embedded in GelMA hydrogel. Scale bar: 50 μm. c) Particle size distribution as measured by NTA. d) Expression of the membrane surface proteins CD9 and TSG101 detected by Western blot analysis.

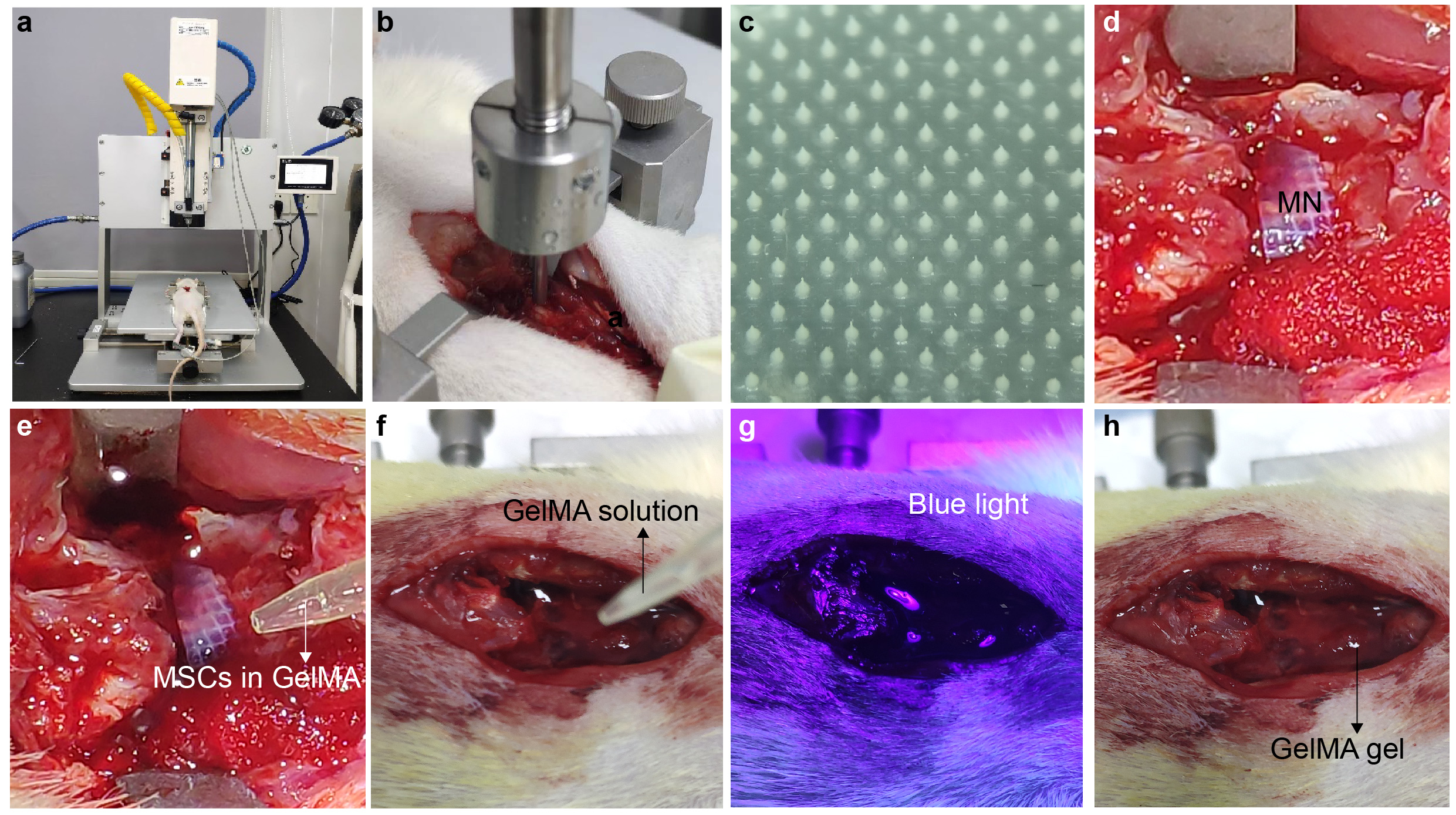

**Figure S2**. The procedures for implanting the MN-MSC patch to treat a rat with SCI. a) Infinite vertical impactor. b) SCI construction . c) Fabricated MN array. d) Implantation of the MN array after dura removal. e) Construction of the MN-MSC patch *in vivo*. f) A GelMA solution was added. g) Blue light crosslinking of GelMA on the SCI site. h) The encapsulation of the MN-MSC patch and SCI with gelled GelMA.

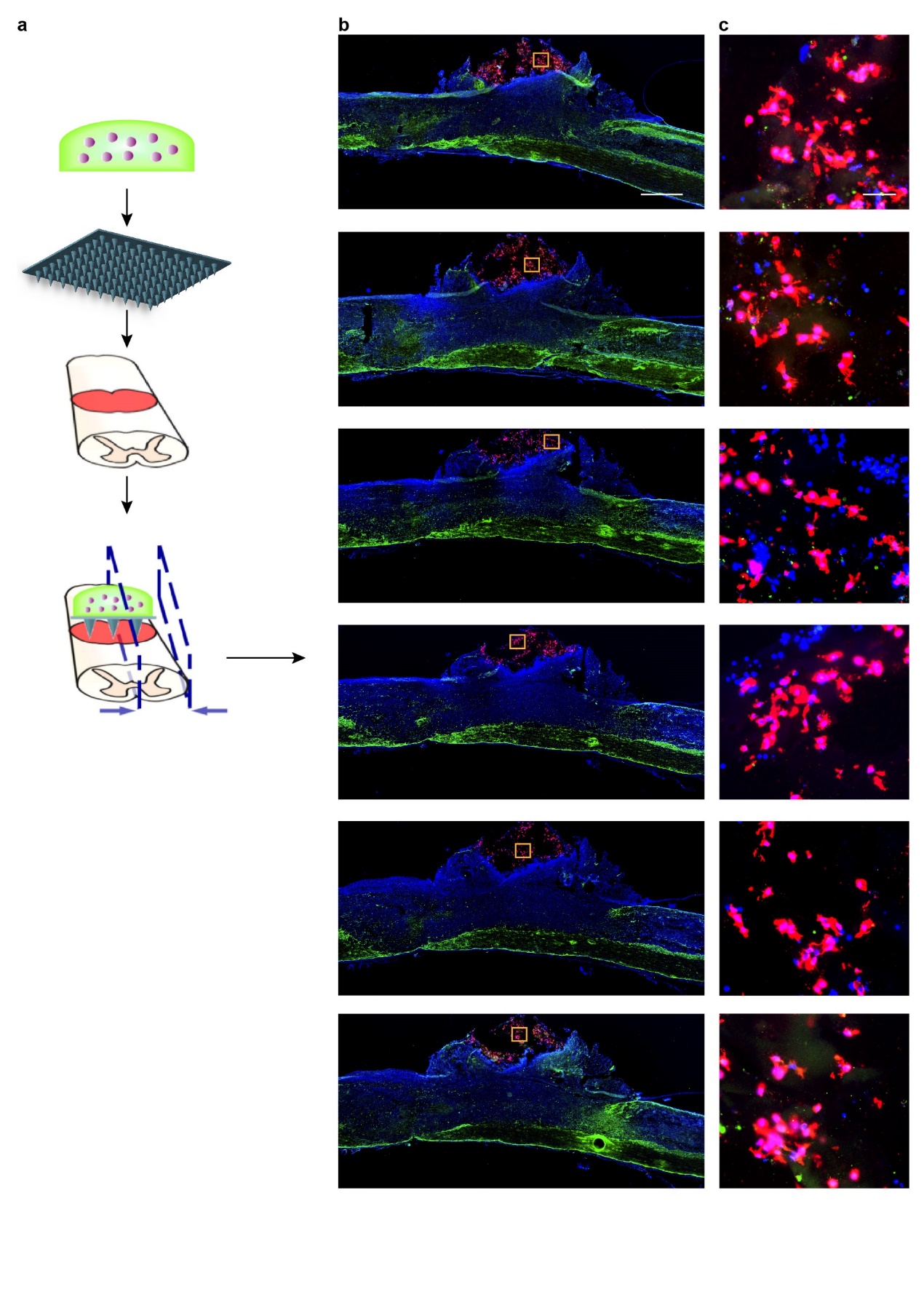

**Figure S3**. Representative serial images of sagittal sections of the spinal cord of a rat treated with MN-MSC patch, green (GFAP), blue (DAPI), and red (GAPDH). a) Schematic illustration of injured spinal cord treated with an MN-MSC patch. b) Serial images of sagittal sections of different positions from the border to the center. Scale bar: 1 mm. c) Show details of MSCs encapsulated in the MN-MSC patch. Scale bar: 50 µm.

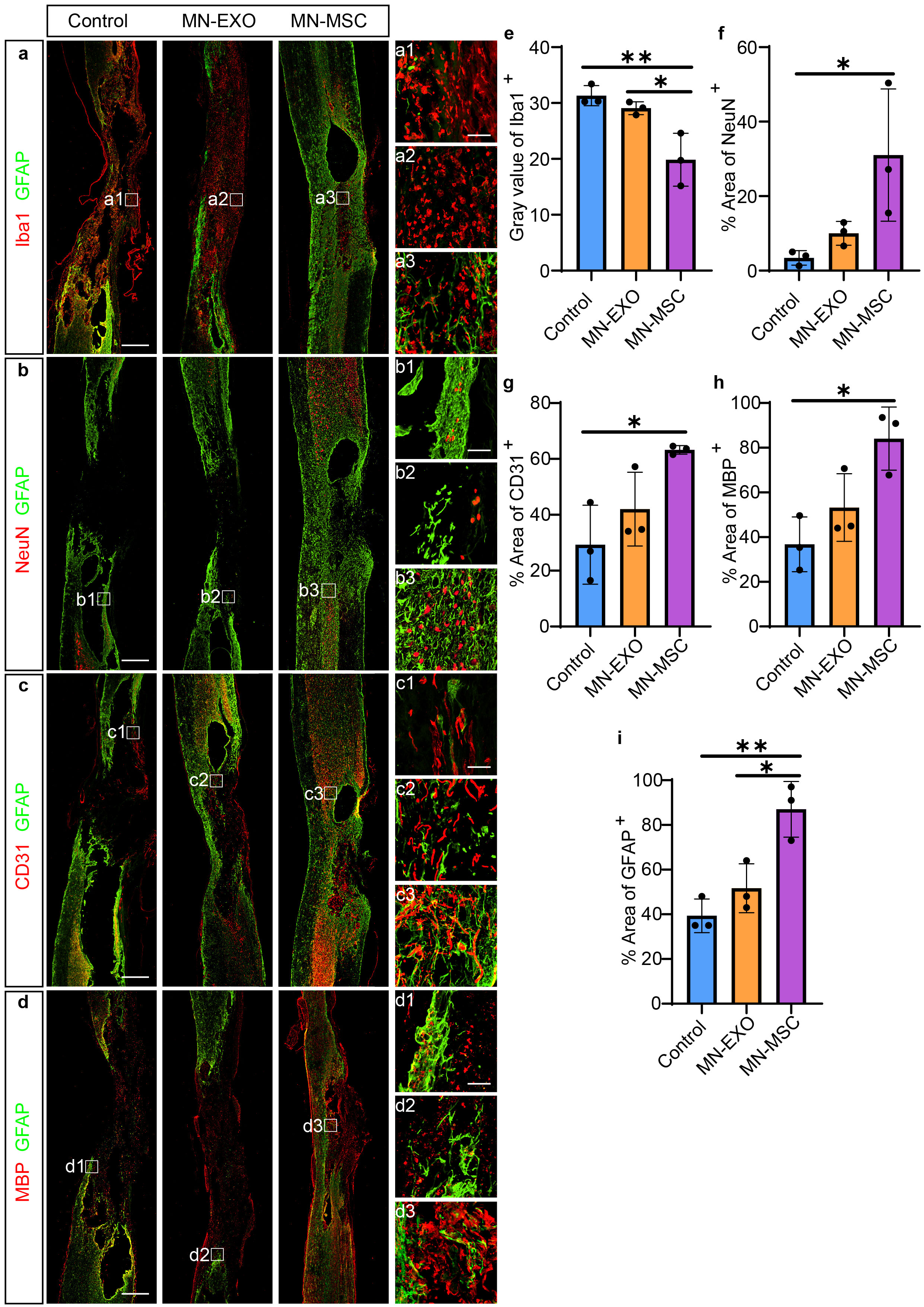

**Figure S4**. Representative images of immunohistochemical staining for GFAP (green), a) Iba1, b) NeuN, c) CD31 and d) MBP (red) of the three groups (control, MN-EXO, and MN-MSC) 8 weeks after injury. Scale bars of image a-d) indicate 1 mm, a1-a3, b1-b3, c1-c3, d1-d3) indicate 100 µm. e-i) Quantification of e) Iba1 immunoreactivity gray values, areas of f) NeuN, g) CD31, h) MBP and GFAP on the injured spinal cord of the three groups. Data are shown as the mean ± SEM. Statistical analysis was performed using one-way ANOVA followed Tukey's multiple comparisons test. n = 3. *p < 0.05, **p < 0.01.

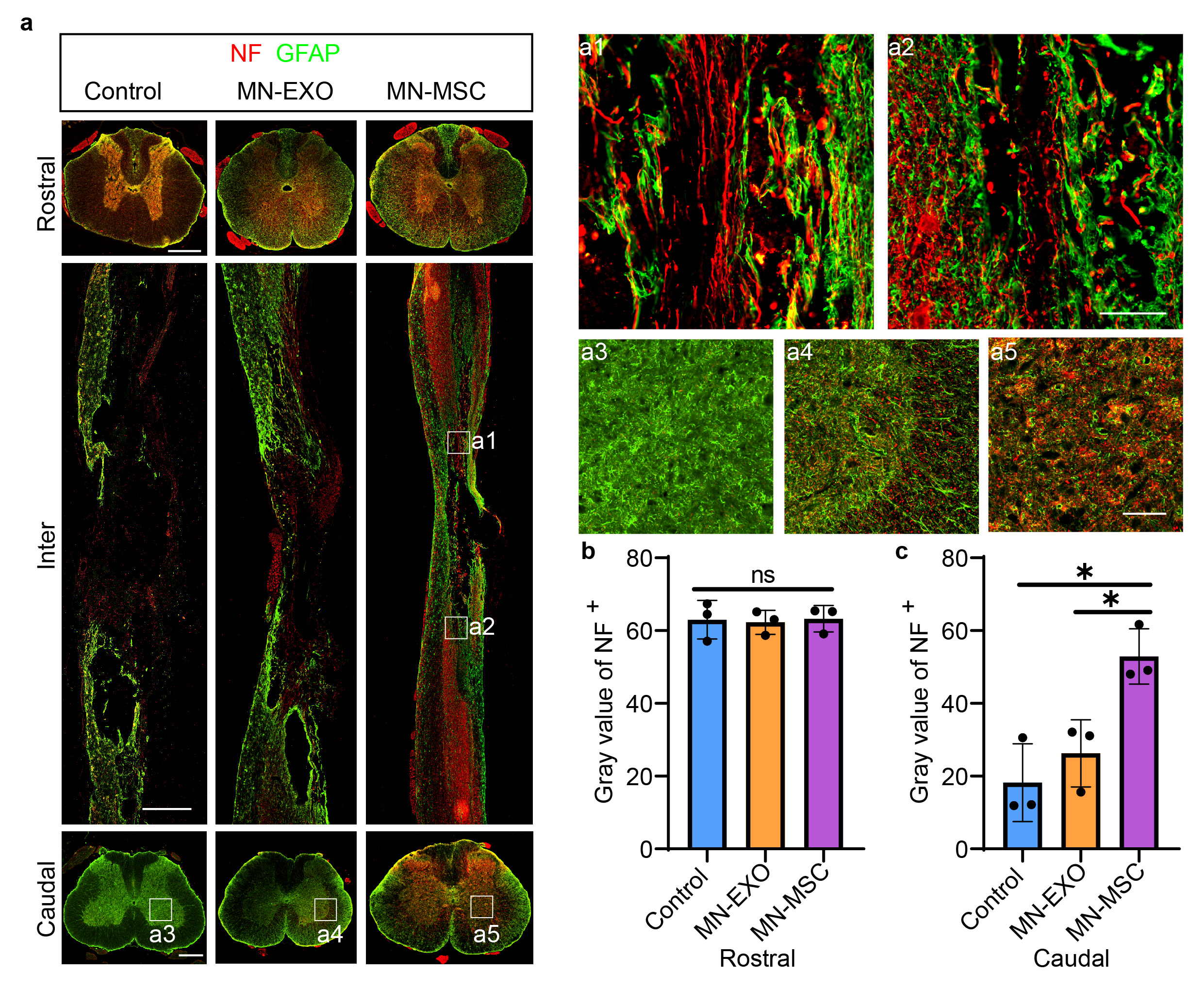

**Figure S5.** a) Representative images of the spinal sections stained with GFAP (green), and NF (red) of rats in the three groups (control, MN-EXO, and MN-MSC). Scale bar: 1 mm. 100 µm. Scale bars of rostral and caudal indicate 500 µm, Inter indicate 1 mm, a1-a2) indicate 100 µm, a3-a5) indicate 100 µm. b-c) Quantification of the NF immunoreactivity gray values on the rostral and caudal sides. Data are shown as the mean ± SEM. Statistical analysis was performed using one-way ANOVA following Tukey’s multiple comparisons test. N = 3. *p < 0.05.

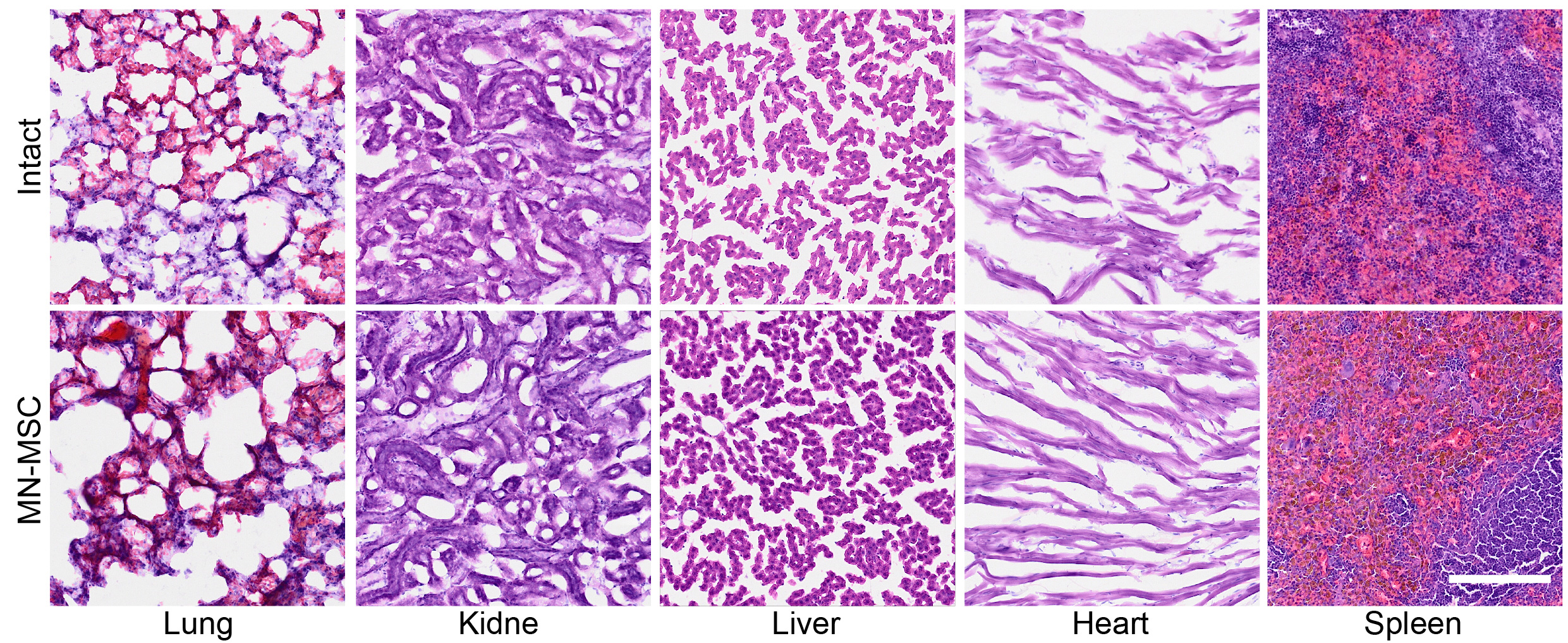
**Figure S6**. H&E staining of the main organs of rats in the intact and MN-MSC groups. Scale bar: 200 μm.

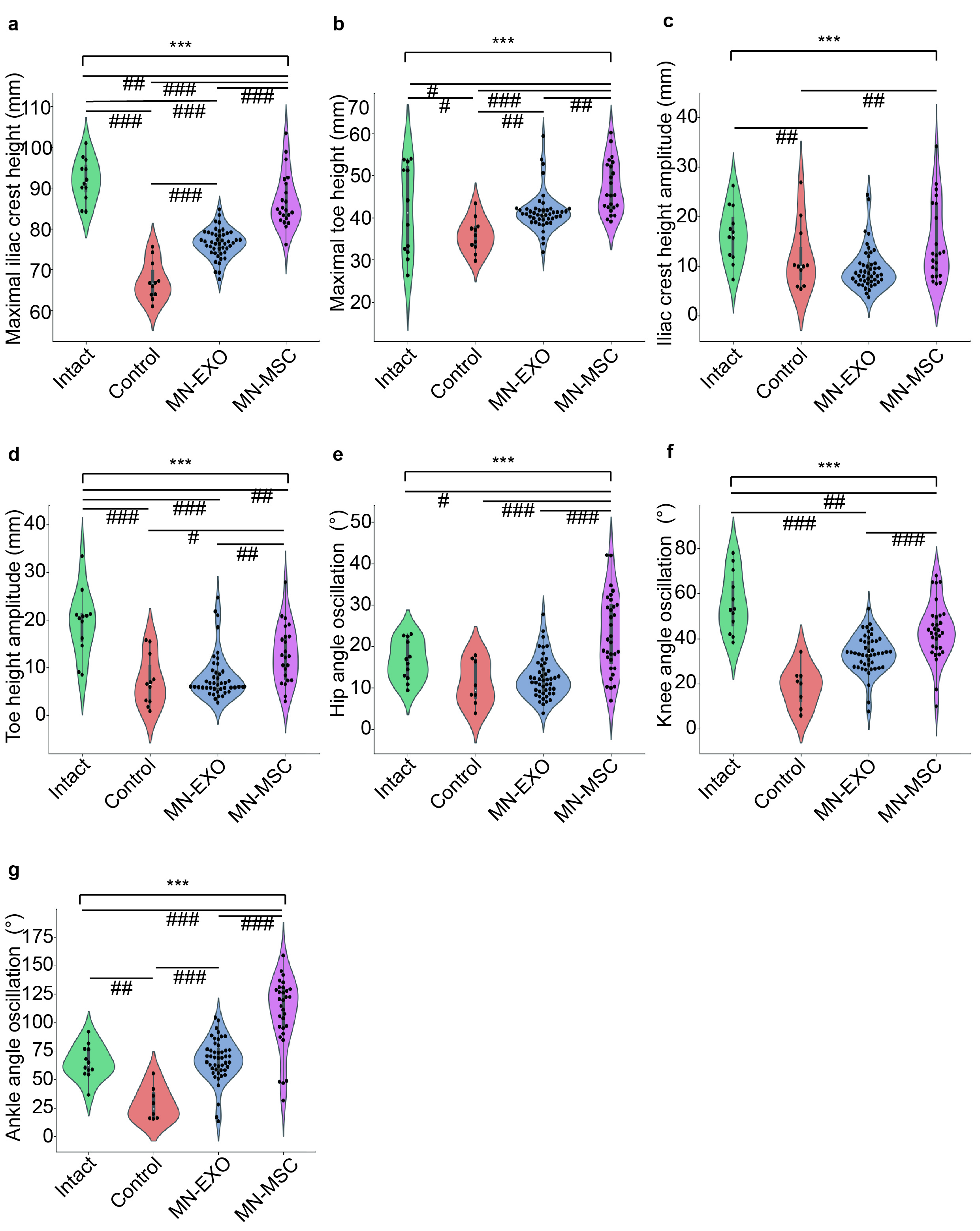

**Figure S7**. Detailed statistical analysis of the hindlimb movement of rats in different groups. Quantification of a) the average maximal iliac crest height, b) maximal toe height, c) iliac crest height amplitude, d) toe height amplitude, e) hip angle oscillation, f) knee angle oscillation, and g) ankle angle oscillation in the different groups. One-way ANOVA with Tukey’s post-hoc test for comparisons among multiple groups (*) and two-tailed paired t tests were used for comparisons within groups (#) for the data shown in the violin plot. n = 8-19. ***p (or ###p) < 0.001, **p (or ##p) < 0.01, *p (or #p) < 0.05.

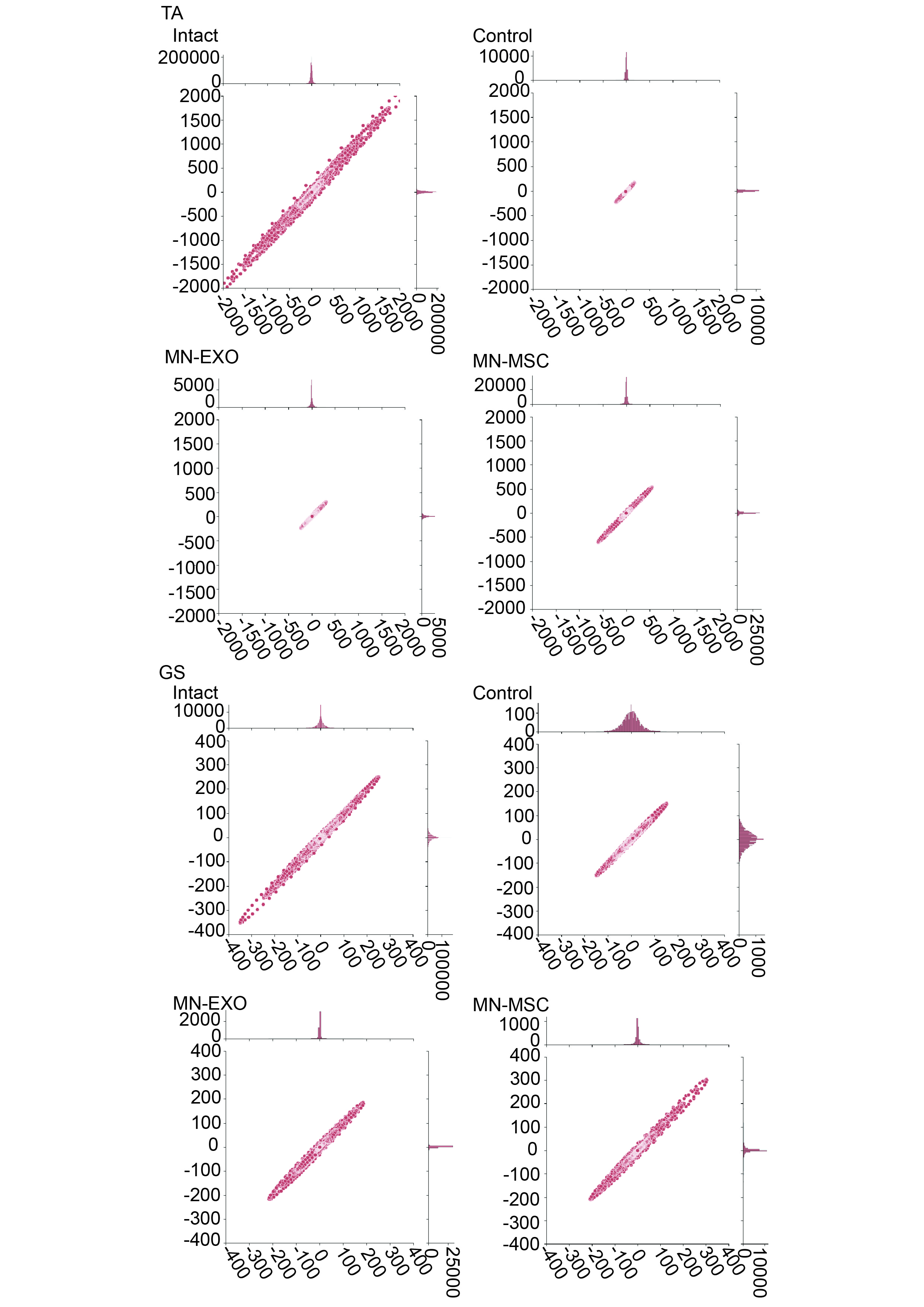

**Figure S8**. Poincaré statistical analysis of the EMG signal amplitude rhythm of TA and GS muscles from rats in the intact, control, MN-EXO and MN-MSC groups.

**Movie S1**. The movements of the hindlimbs of rats under 4 conditions (intact, control, MN-EXO, and MN-MSC).
